# DoUble resin casting micro computed tomography (DUCT) reveals biliary and vascular pathology in a mouse model of Alagille syndrome

**DOI:** 10.1101/848481

**Authors:** Simona Hankeova, Jakub Salplachta, Tomas Zikmund, Michaela Kavkova, Noemi Van Hul, Adam Brinek, Veronika Smekalova, Jakub Laznovsky, Josef Jaros, Vitezslav Bryja, Urban Lendahl, Ewa Ellis, Edouard Hannezo, Jozef Kaiser, Emma R Andersson

## Abstract

**BACKGROUND AND AIMS:** Alagille syndrome, like several other liver diseases, is characterized by malformation of lumenized structures, such as the circulatory or biliary systems. Liver architecture has typically been studied through 2D sections and, more recently, using thick tissue sections combined with immunofluorescence. We aimed to develop a robust method to image, digitalize and quantify 3D architecture of the biliary and vascular systems in tandem.

**METHODS:** The biliary and portal vein trees of the mouse liver were injected with Microfil resin, followed by microCT scanning. Tomographic data was segmented and analyzed using a MATLAB script we wrote to investigate length, volume, tortuosity, branching, and the relation between the vascular and biliary systems. <u>D</u>o<u>u</u>ble resin <u>c</u>asting micro computed <u>t</u>omography (DUCT) was applied to a mouse model for Alagille syndrome (*Jag1*^*Ndr/Ndr*^ mice), in which the biliary system is absent at postnatal stages, but regenerates by adulthood. Phenotypes discovered using DUCT were validated with cumbersome consecutive liver sections from mouse and human liver including patients with Alagille syndrome.

**RESULTS:** DUCT revealed tortuous bile ducts either placed further from portal veins, or ectopically traversing the parenchyma and connecting two portal areas, in *Jag1*^*Ndr/Ndr*^ mice. Furthermore, bile ducts either ended abruptly, or branched independently of portal vein branching, with bifurcations placed hilar or peripheral to portal vein branches. The branching defects, parenchymal bile ducts, and blunt endings were confirmed in patient samples.

**CONCLUSION:** DUCT is a powerful technique, which provides computerized 3D reconstruction of casted networks. It exposes and quantifies previously unknown vascular and biliary phenotypes in mouse models, revealing new phenotypes in patients.

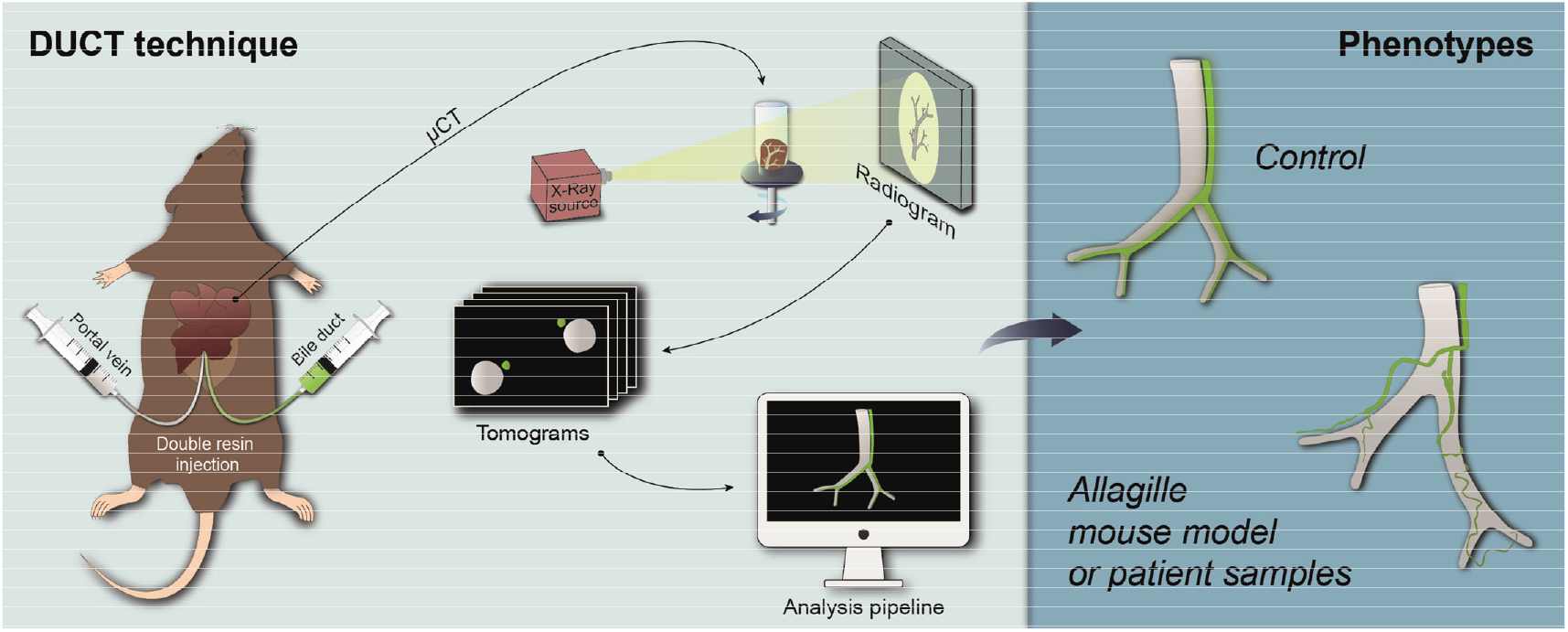

## INTRODUCTION

The three-dimensional (3D) architecture of lumenized structures in our bodies is essential for function and health. The cardiovascular system, lungs, kidneys, liver and other organs depend on precisely patterned tubular networks. Vascular architecture defects contribute to Alzheimer’s disease ^1^; opportunistic infections cause narrowing of bile ducts in liver ^2^; and branching morphogenesis defects in the renal urinary system cause hypertension ^3^. In some pathologies, several lumenized systems are simultaneously affected. Visualizing multiple tubular structures in tandem, in animal disease models, is necessary to allow investigation of how these systems interact in vivo in development, in homeostasis, and in disease. Until now, imaging and quantifying 3D architecture, of multiple structures at once, has been a major challenge in the field.

During embryonic development of the liver, the portal venous system acts as a scaffold and instructional cue for bile duct development, inducing differentiation of adjacent hepatoblasts into cholangiocytes, which undergo tube formation to form ducts (for review see ^4^). Development of the hepatic artery completes the functional unit known as the portal triad. After birth, these three structures show a degree of interdependency. For example, hepatic artery ligation results in bile duct breakdown ^5^. The interplay of these three structures in development and disease is thus crucially important, yet we lack tools to easily visualize and quantify the architecture of these networks in animal models.

Alagille syndrome (ALGS) is a congenital disorder affecting multiple organs, including the hepatic and cardiovascular systems ^6,7^. The disease is usually caused by mutations in the Notch ligand *JAGGED1* (*JAG1*, OMIM: ALGS1 ^8–10^) or, more infrequently, in the receptor *NOTCH2* (OMIM: ALGS2 ^6,11^). Cardiovascular abnormalities include both cardiac malformations such as Tetralogy of Fallot, pulmonary stenosis or septation defects as well as vascular anomalies leading to an increased risk of spontaneous bleeds 12. Hepatic defects are chiefly characterized by peripheral bile duct paucity, though some patients are reported to spontaneously recover a biliary system ^13^, probably via Notch-independent hepatocyte trans-differentiation ^14^.

There are a number of mouse models for ALGS ^15–17^, which recapitulate important aspects of ALGS (reviewed in ^18^). We recently developed a model for ALGS, in a mixed C3H/C57bl6 genetic background, harboring a single nucleotide substitution resulting in a H268Q amino acid change in EGF-like repeat 2 of Jagged1 ^19^. The mouse model is nick-named Nodder (Ndr, *Jag1*^*Ndr*^), due to head-nodding behavior in the heterozygous mice. *Jag1*^*Ndr/Ndr*^ mice recapitulate the cardiac, hepatic (bile duct paucity), ocular, and craniofacial dysmorphology seen in patients. Importantly, this mouse model, also spontaneously recovers a biliary system in the adult stage ^17^.

In order to fully analyze liver vasculopathy or biliary dysmorphology in mouse models, we developed a robust pipeline to visualize and quantify the 3D architecture of the intrahepatic biliary and portal vein systems. Building on extensive meticulous previous work to image intrahepatic architecture using resin ^20–22^, or ink ^23^, we devised <u>d</u>ouble resin-casting micro (<u>μ</u>) computed tomography (DUCT) to image and analyze two networks in 3D, in tandem. To our knowledge, this is the first tool that digitally reconstructs both structures in 3D simultaneously, allowing an in-depth understanding of their relationship. We provide a Matlab pipeline, deposited in https://github.com/JakubSalplachta/DUCT to facilitate adoption of the technique. We applied DUCT to the *Jag1*^*Ndr/Ndr*^ mouse model for ALGS, discovering several unreported phenotypes, demonstrating the strength of this technique to reveal unprecedented pathologies as well as physiological stereotype structures. These phenotypes were compared to tedious consecutive sections in mouse liver and prompted discovery and validation of these phenotypes in liver samples from patients with ALGS.

## MATERIALS AND METHODS

### Experimental mice

All animal experiments were performed in accordance with Stockholms Norra Djurförsöksetiska nämnd (Stockholm animal research ethics board, ethics approval numbers: N150/14 and N61/16) regulations. Animals were maintained with standard day/night cycles, provided with food and water ad libitum, and were housed in cages with enrichment. Adult animals between 4.5 – 6.6 months (both males and females) old were used. *Jag1*^*Ndr/+*^ (Nodder) mice were bred and genotyped as previously described ^22^.

### Patient samples

Collection of liver samples from patients or donors was approved by the Swedish Ethical Review Authority (2017/269-31). Patient samples were obtained at time of liver transplant from extirpated liver, or from left-over donor material, or from organ donation post-mortem. Samples were dissociated for primary cell culture eg. organoids (data not shown), and a matching sample was fixed or fresh-frozen for comparative analyses. Samples from four patients with Alagille syndrome were analyzed as well as two healthy controls.

### MICROFIL® injections

MICROFIL® (Flow Tech Inc.) resin was injected post mortem into common bile duct and portal vein of mouse liver subsequently following transcardial perfusion with HBSS. The liver was dissected out and the resin hardened overnight (ON). The liver was fixed by 3.7% formaldehyde and separated into lobes. Right medial lobe was used for micro computed tomography (microCT) scanning. For details, see the Supplementary Material.

### Whole mount immunohistochemistry

Mice were anesthetized and transcardially perfused with HBSS and 10% neutral buffered formalin (NBF) for 5 min. Liver was dissected out and further immersion fixed with 10% NBF ON at 4°C. The next day liver was washed and kept in DPBS and separated into lobes. The right medial lobe was stained, cleared following the iDISCO+ protocol and imaged by light sheet microscope (performed by Gubra (Denmark). For details, see the Supplementary Material.

### Ink Injections

Ink (Higgins cat. #44032) was injected post mortem into common bile duct and portal vein of mouse liver subsequently following transcardial perfusion with HBSS. The liver was dissected out, fixed by 3.7% formaldehyde and separated into lobes. Right medial lobe was used for clearing with Benzyl Benzoate and Benzyl Alcohol and imaged with stereomicroscope Stemi 305 (Carl Zeiss Microscopy) camera (Cannon, PowerShot S3 IS). For details, see the Supplementary Material.

### Liver immunohistochemistry

Adult mouse livers were collected either fresh or fixed ON in 3.7% FA. Human liver was fixed with 3.7% FA. Five μm formalin-fixed, paraffin-embedded constitutive liver sections were utilized for chromogenic staining. For immunofluorescence staining fourteen μm fresh frozen cryosections were used. For detailed protocol, see the Supplementary Material.

### Image processing

Whole mount liver iDISCO+ cleared images were initially processed in ImageJ (NIH) and further filtered and analyzed in Amira (Thermo Fisher Scientific). Ink injected bile duct and portal vein filament tracing was performed on 2D images using Amira. Liver section images were adjusted for contrast, brightest and fluorescent levels in ImageJ. For more details see the Supplementary Material.

### Micro CT measurement

The system GE Phoenix v|tome|x L 240 (GE Sensing & Inspection Technologies GmbH, Germany) equipped with nanofocus X-ray tube (180 kV/15 W) was used for the tomographic measurements that were carried out in the air-conditioned cabinet (fixed temperature 21°C). To prevent any sample motion during the scanning, the samples were placed in 15 ml Falcon tube, filled with 1% agarose gel. The tomographic reconstruction of acquired data was performed using GE phoenix datos|x 2.0 software. The voxel resolution was fixed for all measurements at 12 μm, except one (sample #2401, 8 μm). Detailed overview of used acquisition parameters is stated in Table 1 in the Supplementary Material.

### Micro CT data segmentation

The resin segmentation was performed by global thresholding (Supp Fig 11A, A’) and supplemented with manual corrections of the resin cast artefacts (e.g. air bubbles or resin leakage due to lumen rupture, Supp Fig 1B) using VG Studio MAX 3.2 (Volume Graphics GmbH) software. 3D interactive. PDF files are provided in the Supplementary Material for the best assessment of canal complexity (Supp Fig 3 – 8). For more details see the Supplementary Material.

### Micro CT data analysis

To analyze morphological parameters such as volume, length, diameter, distance, tortuosity and branching of the BD and PV systems using segmented 3D binary masks, a custom-written algorithm and freely available Matlab® codes (Version R2017a, The MathWorks Inc., Natick, MA) were developed. The algorithms are based on skeletonization of the input binary masks (using homotopic thinning algorithm described in ^24^) and conversion of calculated 3D medial axis skeleton to a network graph (using algorithm described in ^25^). For more information and implementation specifics see the Supplementary Material.

### Statistical analysis

The control measurements were compared to *Jag1*^*Ndr/Ndr*^ data using Student’s *t*-test (Fig 2B – 2D, 3D – 3J, 5C and Supp Fig 9B – 9K) or 2-way ANOVA (Fig 2E – 2F, 4B, 4D, 4E, 5B and Supp. Fig 9L, 9M) followed by multiple comparison. *p* value bellow 0.05 was considered statistically significant.

## RESULTS

### DUCT outperforms other state of the art techniques to visualize liver in 3D

Current state of the art methods to visualize 3D architecture include (1) ink injection followed by organ clearing, (2) corrosion casting and tissue maceration, and (3) whole mount immunofluorescence or fluorescent reporter imaging in cleared organs. Ink injection (Fig 1 A – F) and corrosion casting are useful for basic assessment of morphology, but analysis and data dissemination are limited by the mode of image acquisition, which is typically photography or scanning electron microscopy ^26^ (both 2D methods). Dual ink injection of the biliary and vascular systems allows visualization of fine blood vessels and ductules (Fig 1B), but their quantification requires separate injection and imaging (Fig 1C, D). 2D quantification of different parameters is achieved by filament tracing in image processing software (Fig 1E, F). Another commonly used method is whole-mount imaging of fluorescently labelled cells, using antibodies or reporters, with organ clearing techniques such as iDISCO. We applied iDISCO+ to a mouse liver with antibody staining for cholangiocytes (CK7) and smooth muscle cells (alpha SMA) (Fig 1G – L). Fixation, antibody quality and permeabilization issues result in highly variable outcomes(Supp Fig 1A). Similar to ink injections, fine architecture of terminal ductules and blood vessels are well resolved in the periphery of the liver (Fig 1H) where penetrance is less of an issue, but not in the bulk of the liver (Fig 1L, Supp Fig 1A). Analysis of 3D structure is standard in whole mount imaging, and software tools including Imaris and Amira can be used to segment the vascular or biliary systems. While this is a powerful approach, diseases in which biliary cells are present but clustered, and the ducts lack a lumen, are difficult to assess using whole mount fluorescence, which focuses on cell markers rather than lumenized structures.

**Fig. 1.**
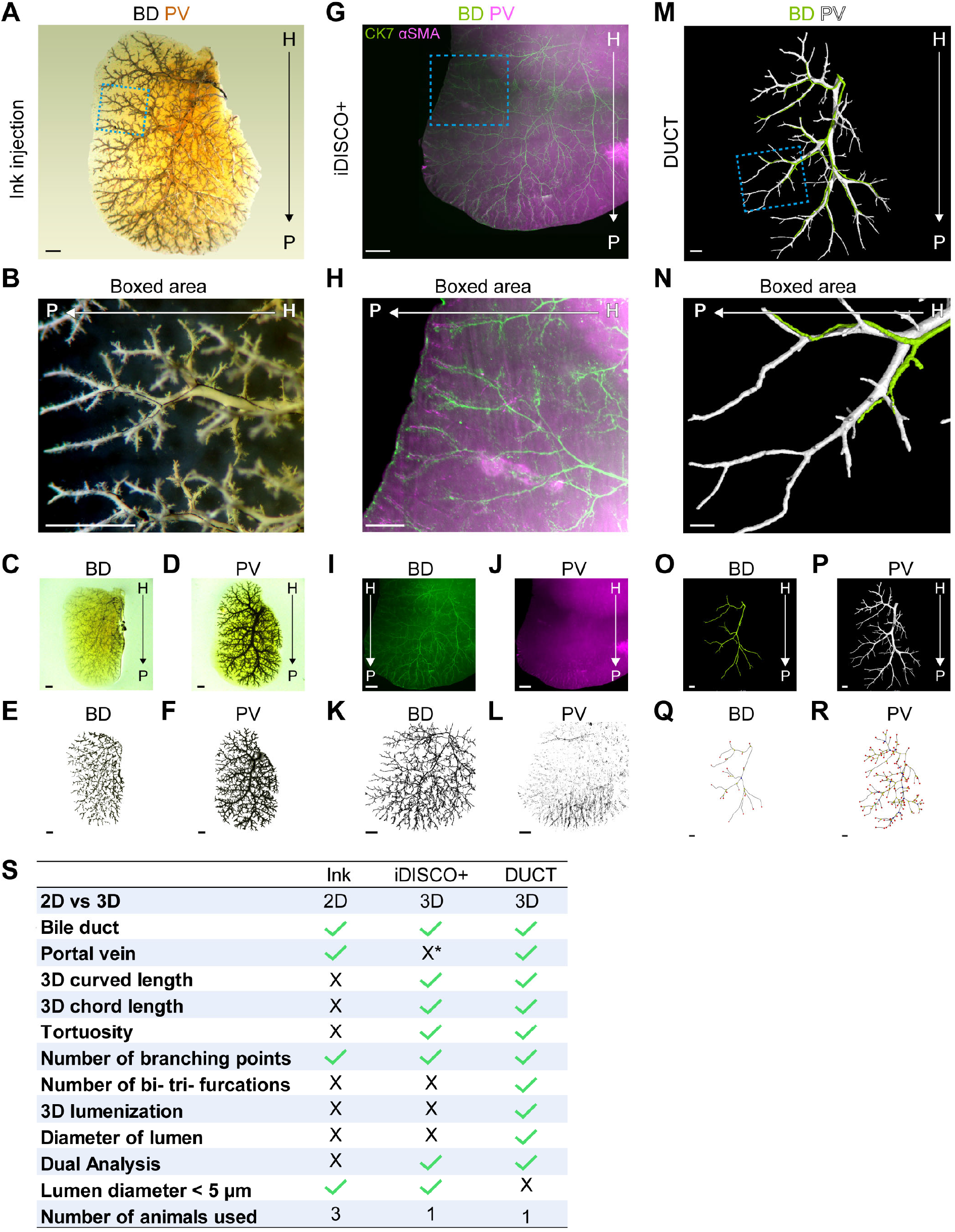
DUCT outperforms other state of the art techniques to visualize liver in 3D. (A – F) Whole liver ink injection into portal vein (PV) and bile duct (BD), followed by clearing, 2D imaging and filament tracing. (G – L) Whole mount immunofluorescence staining coupled with iDISCO+ clearing and filament tracing. (M – R) Double resin injection into a portal vein and bile duct scanned with micro computed tomography (CT) and combined with skeletonization and identification of branching nodes. (S) Table compares the three techniques in means of dimensionality, measurements and 3Rs. * Refers to our results, in principle portal vein can be visualized. Scale bars (A, C – F, G, I – L, M, O – R) 1 mm, (B, H, N) 500 μm.

To address the 3D architecture of different tubular systems, resin casting followed by microCT has emerged as a powerful technique to obtain volumetric and branching data ^20,22,27–29^. However, this approach has not yet been multiplexed to simultaneously assess several lumenized structures at once. We therefore injected the biliary and portal vascular systems with two different resins (MICROFIL®) and subjected the liver lobe to microCT. We dubbed the combination of double injection with the customized Matlab pipeline double resin casting micro (<u>μ</u>) computed tomography (DUCT) (Fig 1 M – R). While each method offers certain advantages, DUCT is the only approach that quantifies the 3D lumenization of the vascular or biliary system simultaneously (Fig 1Q, R, S). Manual segmentation of the two systems assigns a portal vein or biliary identity to the casts, and casts are inspected for deficient perfusion and air bubbles (Supp Fig 1B), which are manually corrected or filled in. Concurrently, livers were cleared to verify perfusion visually (Supp Fig 1C). Poorly perfused livers were not included in the analysis (Supp Fig 1D). The Matlab pipeline then quantifies the curved length (actual length), chord length (theoretical length, assuming shortest distance between two points), diameter, volume, bifurcations and trifurcations of the vascular and biliary systems, and the relation between them such as whether bile ducts (BDs) branch adjacent to portal vein (PV) branches and the distance between BD and nearest PV. A DUCT limitation is the restricted lumen size (max 5 μm) that the resin can penetrate, thus prohibiting analysis of the finest branches of the biliary or vascular trees (Fig 1S). However, DUCT exceeded the ink injections and iDISCO+ whole mount immunofluorescent staining in all other quantifiable parameters, as well as in dual analysis, while reducing the number of animals used per experiment.

### Jag1^Ndr/Ndr^ bile ducts regenerate to form a fully-grown biliary network

To test DUCT, we applied the method to a mouse model for ALGS, at adult stages in which bile ducts have regenerated. We employed DUCT to *Jag1*^*Ndr/Ndr*^ mice to investigate (1) the architecture of the regenerated biliary system, (2) whether the intrahepatic vascular system is affected and (3) how these two systems spatially interact in normal and diseased liver. We assessed the architecture in the right mediolateral lobe, in order to use other lobes for quality control and immunohistochemistry. All *Jag1*^*Ndr/Ndr*^ adult livers displayed lumenized bile ducts (Fig 2A, Supp Fig 2). While *Jag1*^+/+^ mice demonstrated a stereotyped vascular and biliary architecture (Supp Fig 2A, B, C), *Jag1*^*Ndr/Ndr*^ livers exhibited greater architectural variability (Supp Fig 2 E – G). MicroCT scans can be explored in separate channels or in tandem (interactive PDFs, Supp Fig 3 – 8). Quantification of the individual features of biliary and vascular structures was performed by first creating a mask to address the branching properties of the PV (Fig 1Q, Supp Fig 2D, G, left panel) and BD (Fig 1R, Supp Fig 2D, G, right panel) with bifurcations labelled in yellow, trifurcations in blue, branches with >3 nodes in green, and branch termination points in red.

**Fig. 2.**
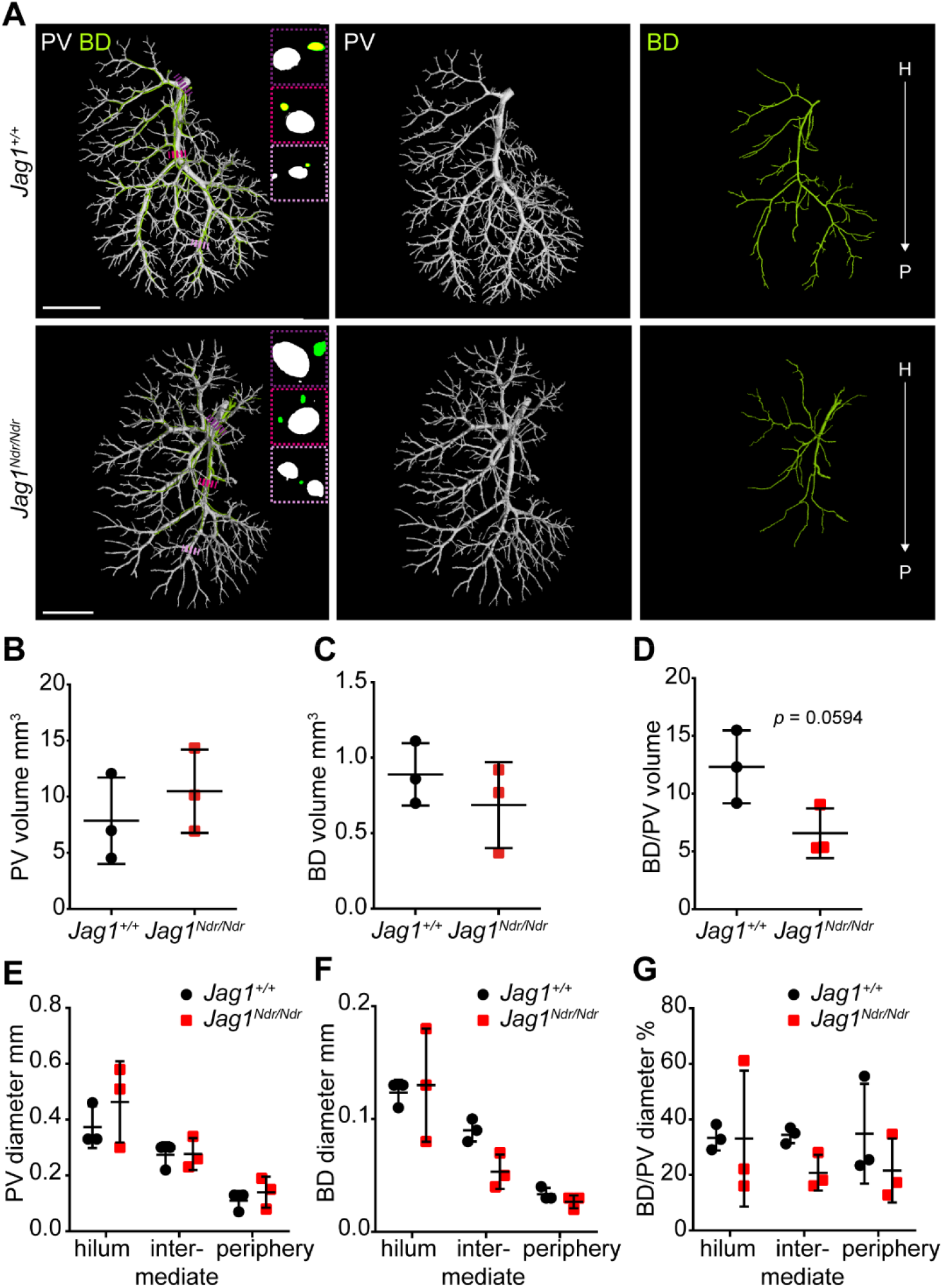
*Jag1*^*Ndr/Ndr*^ bile ducts regenerate to form a fully-grown biliary network. (A) 3D rendering of BD and PV structures using DUCT in *Jag1*^+/+^ (top panel) and *Jag1*^*Ndr/Ndr*^ (bottom panel). Boxed regions highlight 2D section through hilar, intermediate and peripheral PV. Scale bar 4 mm. (B) Overall volume of the portal vein system. (C) Overall volume of the biliary system. (D) BD to PV system volume ratio. (E, F) PV and BD diameter analysis. (G) BD to PV diameter ratio. (B – G) Each dot represents one animal, lines show mean value ± standard deviation. Statistical test (B – D) unpaired *t*-test, (E – G) two way ANOVA.

The total volume of the vascular and biliary system was similar in *Jag1*^*Ndr/Ndr*^ and *Jag1*^+/+^ mice (Fig 2B, C). However, there was a tendency towards a larger PV volume and smaller BD volume in *Jag1*^*Ndr/Ndr*^ mice, resulting in a slightly skewed BD/PV volume ratio (*p* = 0.0594). Next, we investigated the PV and regions) There were no consistent differences between *Jag1*^+/+^ and *Jag1*^*Ndr/Ndr*^ mice (Fig 2E, F), and *Jag1*^*Ndr/Ndr*^ mice displayed high variability in diameter in the hilar region (Fig 2E, F). The BD to PV diameter ratio was ca 1:3 in hilar and intermediate regions of *Jag1*^+/+^ liver (BDs have 30% the diameter that PVs have) (Fig 2G). The BD/PV diameter ratio was not preserved in the *Jag1*^*Ndr/Ndr*^ mice (Fig 2G). In conclusion, the *Jag1*^*Ndr/Ndr*^ biliary system regenerated with minor shifts towards smaller volume and diameter, while the portal vasculature increased in these parameters.

### Jag1^Ndr/Ndr^ liver regrows postnatal tortuous bile ducts

Next, we investigated the length and tortuosity of the biliary and portal vascular trees. 3D reconstruction of PV vasculature and the biliary network revealed straight *Jag1*^+/+^ BDs (Fig 3A top panel), whereas *Jag1*^*Ndr/Ndr*^ BDs were tortuous (Fig 3A bottom panel), especially in the liver periphery. We confirmed in serial histological liver sections that the *Jag1*^*Ndr/Ndr*^ BD were tortuous (Fig 3B). Furthermore, we applied the DUCT Matlab pipeline to quantify the actual (curved) length and theoretical (chord) length (scheme Fig 3C). The curved and chord lengths of the entire PV system, the main PV branch alone (Fig 3D, E, Supp Fig 9A, B) the entire BD network, or the main BD branch alone (Fig 3F, G, Supp Fig 9C) did not differ between *Jag1*^+/+^ and *Jag1*^*Ndr/Ndr*^ animals. The ratio between BD and PV curved (Supp Fig 9D) or chord (Supp Fig 9E) was also similar. There was a 6% increase in portal vein tortuosity (Fig 3H, 16% ± 0.14% vs 18% ± 0.23%), while biliary tortuosity was increased by 50% in *Jag1*^*Ndr/Ndr*^ livers (Fig 3I, 20%± 0.91% vs 29% ± 5.07%). The tortuosity of the main branch alone, for PV or BD, was not different (Supp Fig 9 F, G). In summary, *Jag1*^*Ndr/Ndr*^ bile ducts recovered volume and length postnatally but displayed altered morphology with a pronounced increase in tortuosity.

**Fig. 3.**
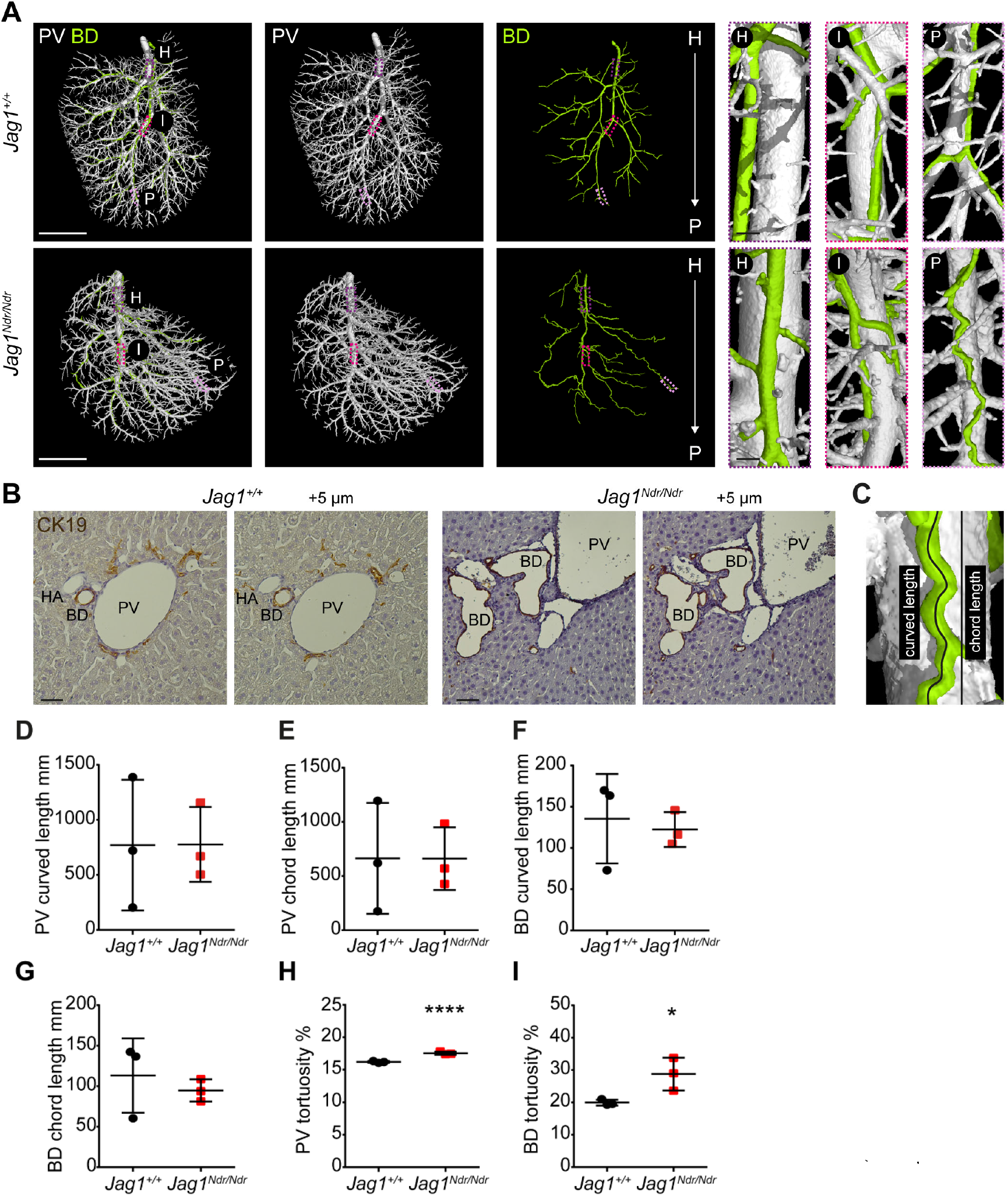
*Jag1*^*Ndr/Ndr*^ liver regenerates postnatal tortuous bile ducts. (A) DUCT 3D rendering of BD and PV structures in *Jag1*^+/+^ (top panel) and *Jag1*^*Ndr/Ndr*^ (bottom panel) liver. Boxed areas represent the hilar (H), intermediate (I) and peripheral (P) regions. Scale bars 4 mm, boxed regions 250 μm. (B) 2D histological consecutive liver sections, scale bar 50 μm. (C) Curved (actual) and chord (theoretical) length measurements scheme. Quantification of curved and chord (D, E) PV lengths and (F, G) BD lengths. (H) PV tortuosity. (I) BD tortuosity. (D – I) Each dot represents one animal, lines show mean value ± standard deviation. Statistical test (D – I) unpaired *t*-test, *p* < 0.05 (*), *p* < 0.0001 (****).

### Regenerated bile ducts in Jag1^Ndr/Ndr^ mice, and in patients with Alagille syndrome, display branching defects

The dual analysis of vascular and biliary systems enabled us to explore the interaction between the two structures. First, we investigated the branching pattern of both trees. During embryonic development, the biliary system is established alongside the PV. This is reflected in the branching morphology of *Jag1*^+/+^ BDs (Fig 4A left panel, blue arrowhead) that branch close to PV branching points (magenta arrowhead). In contrast, *Jag1*^*Ndr/Ndr*^ BDs (Fig 4A middle panel, blue arrowhead) branch further from the PV (magenta arrowhead), or independently of PV branching (Fig 4A right panel, blue arrow; defined as distance > 0.54 mm). Quantification of distance between BD branching point and the nearest PV branching point confirmed significant increase in *Jag1*^*Ndr/Ndr*^ liver (Fig 4B). We analysed consecutive histological liver sections to compare branching morphology. *Jag1*^+/+^ BDs (Fig 4C top panel, blue arrowhead) indeed branch at the same point as PVs branch (magenta arrowhead). In contrast, in *Jag1*^*Ndr/Ndr*^ liver, PVs (Fig 4C middle panel, magenta arrowhead) branch, without BD branching (blue arrowhead), or BDs bifurcate in the absence of PV branching (Fig 4C bottom panel, blue arrow). Utilizing the DUCT Matlab pipeline we evaluated branching (Supp Fig 9 H – M) by quantifying the number of vascular or biliary bifurcations, trifurcations, or nodes with more than 3 branches, but did not identify any difference between *Jag1*^+/+^ and *Jag1*^*Ndr/Ndr*^ systems. Branching trees in biological systems have a stereotype structure in which branch lengths shorten with each branching generation from start (hilum) to end (periphery) ^30^. Using ImageJ we measured the branch length of PV segments and BD segments for eleven generations and found that PV segments shorten as expected with each generation in both *Jag1*^+/+^ and *Jag1*^*Ndr/Ndr*^ livers (grouped data in Fig 4D, per generation in Supp Fig 9O). *Jag1*^+/+^ BDs followed the same stereotype branching principle (grouped data in Fig 4E, per generation in Supp Fig 9P), while *Jag1*^*Ndr/Ndr*^ BD branch length was uniform across the hierarchy of branches and did not decrease between branching generations. Importantly, this indicates that hilar BD segments were shorter than expected and suggest ectopic, regenerative, branching occurs in the hilar region. We next asked whether similar branching phenotypes are present in healthy human liver or in patients with ALGS. In normal liver, BD branching occurred within 35 μm of PV bifurcation (Fig 4F top panel, blue arrowheads). In a liver from a patient with ALGS, with regenerated BDs, we observed PV branching in the absence of BD branching (> 100 μm to nearest branchpoint) (Fig 4F middle panel, blue arrowhead), as well as BDs merging (Fig 4F bottom panel, blue arrows). Regenerating BDs in ALGS thus displayed similar branching abnormalities as described above for the *Jag1*^*Ndr/Ndr*^ liver.

**Fig. 4.**
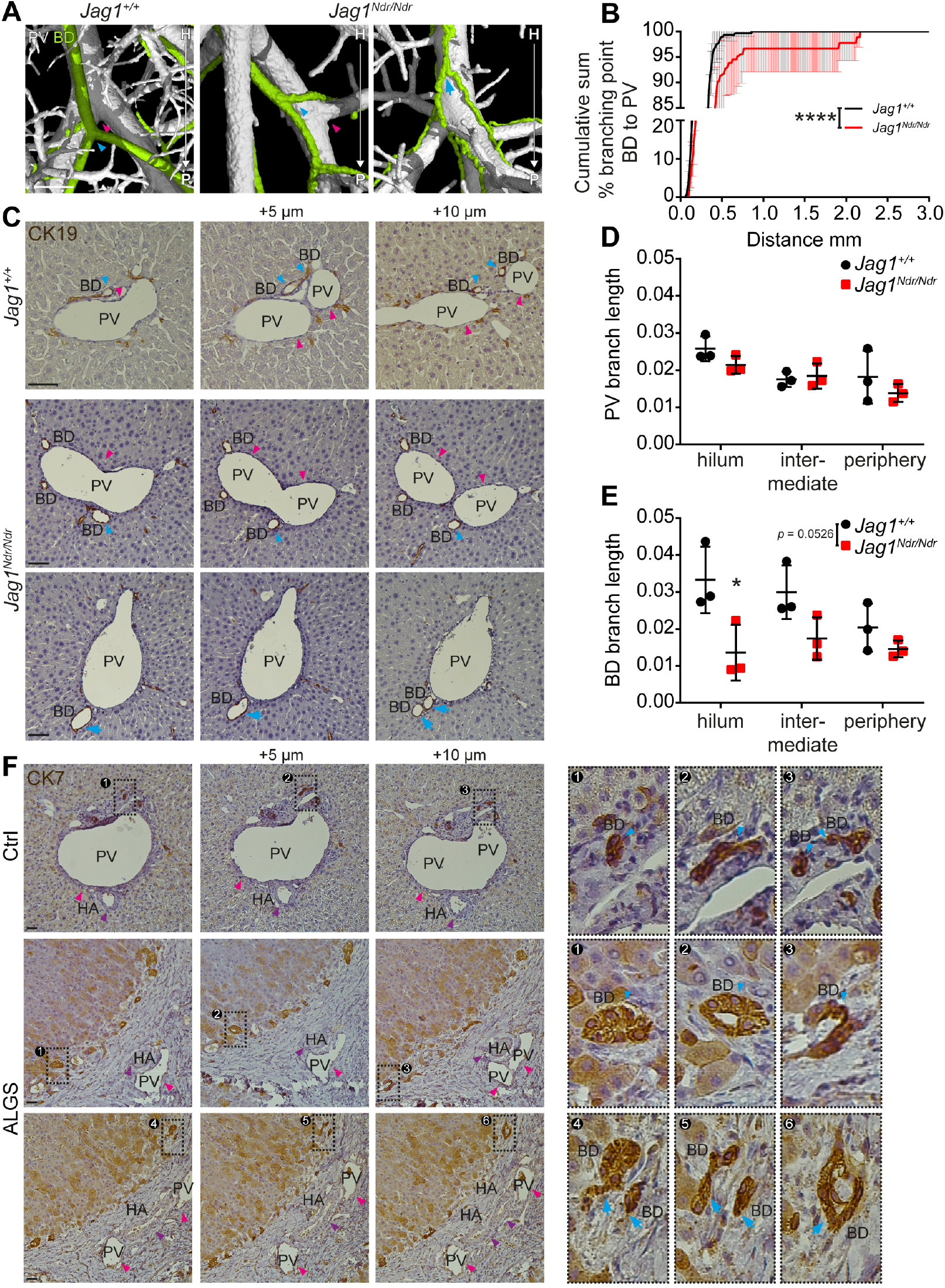
Regenerated bile ducts in *Jag1*^*Ndr/Ndr*^ mice, and in patients with Alagille syndrome, display branching defects. (A) Branching pattern in *Jag1*^+/+^ (left panel) and *Jag1*^*Ndr/Ndr*^ liver (middle and right panel). Scale bar 500 μm. (B) Cumulative sum of % BD branching point at given distance to PV branching point. Middle line represents mean value ± standard deviation. (C) Branching analysis in 2D histological consecutive liver sections, scale bar 50 μm. (D) Analysis of PV and (E) BD branch length. Each dot represents one animal, lines show mean value ± standard deviation. (B, D, E) two-way ANOVA, followed by multiple comparison, *p* < 0.05 (*), *p* < 0.0001 (****). (F) Branching pattern in consecutive human liver histological sections, scale bar 50 μm.

### Regenerated bile ducts are situated further from portal veins in Jag1^Ndr/Ndr^ mice and in patients with Alagille syndrome

During embryonic development, cholangiocytes differentiate from hepatoblasts that are in contact with portal vein mesenchyme expressing *Jag1* ^4^, a process that is disrupted in *Jag1*^*Ndr/Ndr*^ liver ^17^. Whether regenerated bile ducts arise adjacent to portal veins, or whether bile ducts that regenerate, potentially through Tgfbeta-driven transdifferentiation ^14^, are less dependent on proximity to portal veins has not yet been explored. We therefore analyzed the distance between biliary and portal vascular systems. *Jag1*^+/+^ BDs maintain a uniform distance to adjacent PVs (Fig 5A top panel, asterisk, 5B, all BDs within 0.5mm of a PV). In contrast, *Jag1*^*Ndr/Ndr*^ BDs do not maintain a uniform distance to the nearest PV (Fig 5A bottom panel, double arrow, 5B, 1.5% of BD measurement points between and 1.26 mm away from a PV) and sometimes traverse the parenchyma between two portal vein branches (Fig 5A bottom right panel, Supp Fig 1C, D empty arrowheads). Both the increased BD-PV distance and parenchymal bile ducts were validated in histological sections, where *Jag1*^+/+^ BDs were in contact with PV mesenchyme (Fig 5D top panel, asterisk). *Jag1*^*Ndr/Ndr*^ BD were confirmed to be present outside of the portal vein mesenchyme area (Fig 5D middle panel, double arrow) or in the liver parenchyma close to the edge of the liver (Fig 5D bottom panel). This phenotype, visualized in 2D sections, may resemble cholangiocyte proliferation rather than a bridging structure, highlighting the importance of 3D imaging. Parenchymal BD were also detected in liver from patients with ALGS (Figure 5E) but not in normal human liver. These results indicate that the *Jag1*^*Ndr/Ndr*^ biliary tree and regenerated BD in patients with ALGS developed through an alternative mechanism in which cholangiocyte progenitors did not require close contact with the portal vein mesenchyme.

**Fig. 5.**
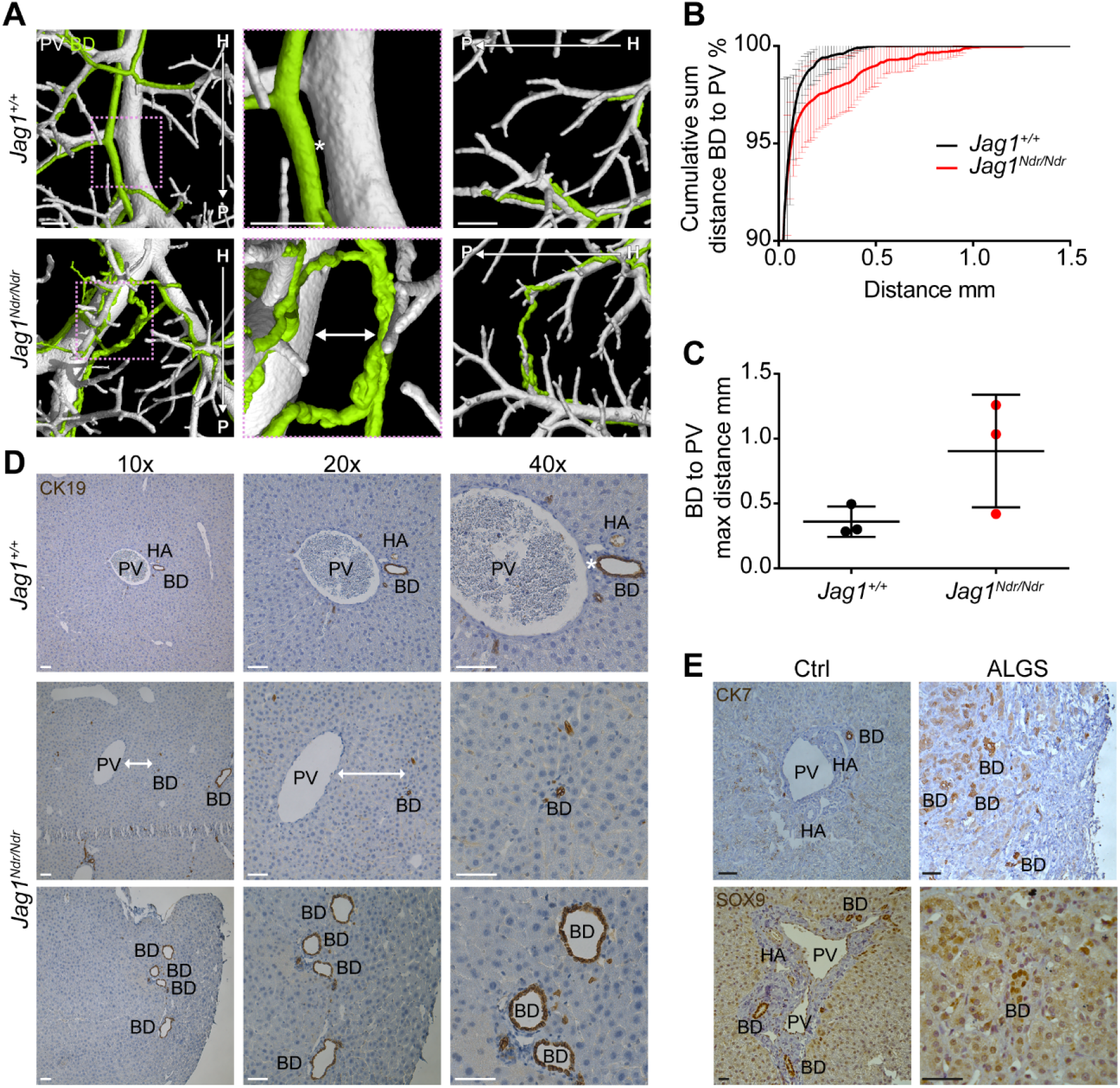
Regenerated bile ducts are situated further from portal veins in *Jag1*^*Ndr/Ndr*^ mice and in patients with Alagille syndrome. (A) Distance analysis between *Jag1*^+/+^ (top panel) and *Jag1*^*Ndr/Ndr*^ (bottom panel) BD and PV trees. Scale bar 500 μm. (B) Cumulative sum of voxel numbers between BD and PV. Middle line represents mean value ± standard deviation, two-way ANOVA. (C) Maximum distance between BD and PV. Each dot represents one animal, lines show mean value ± standard deviation, unpaired *t*-test. (D) Distance analysis in 2D histological sections. Scale bar 50μm. (E) Parenchymal BDs in histological liver sections from patients with ALGS. Scale bar 50μm.

### Bile ducts in Jag1^Ndr/Ndr^ mice and patients with Alagille syndrome terminate abruptly

In mammals, the biliary tree forms by a tubulogenic process in which a heterogeneous, hierarchical fine network composed of connected cholangiocytes is refined to a single larger conduit of bile ducts ^4^. The remaining cholangiocytes that were not incorporated into functional ducts are removed by a mechanism that is still poorly understood ^4^. DUCT enables the visualization of perfused duct canals with a diameter greater than five μm and can thus reveal the connectivity of the biliary network. In *Jag1*^+/+^ liver the biliary system formed a continuous uninterrupted tree (Fig 6A left panel, Supp Fig 2B). The *Jag1*^*Ndr/Ndr*^ biliary system instead displayed branches that appeared to orient from peripheral to hilar, rather than hilar to peripheral, with blunt abrupt endings (Fig 6A right panels, blue arrowhead) (on average one blunt end BD per one lobe). We confirmed the connectivity of BDs in liver histological serial sections and identified continuous BDs in *Jag1*^+/+^ livers (Fig 6B top panel), but abruptly ending BDs in *Jag1*^*Ndr/Ndr*^ livers (Fig 6B bottom panel). The black arrowhead marks a well-formed BD that suddenly ended in the following sections (blue arrowheads) and eventually was visible only as one cytokeratin 19 (CK19) positive cell. Similarly, there were abruptly ending BDs in liver samples from patients with ALGS (Figure 6C). Black arrowhead labels a well-formed BD that terminates in the subsequent section (blue arrowhead). In conclusion, the *Jag1*^*Ndr/Ndr*^ and ALGS regenerated biliary system displays abruptly ending BDs, which may affect bile flow and shear stress.

**Fig 6.**
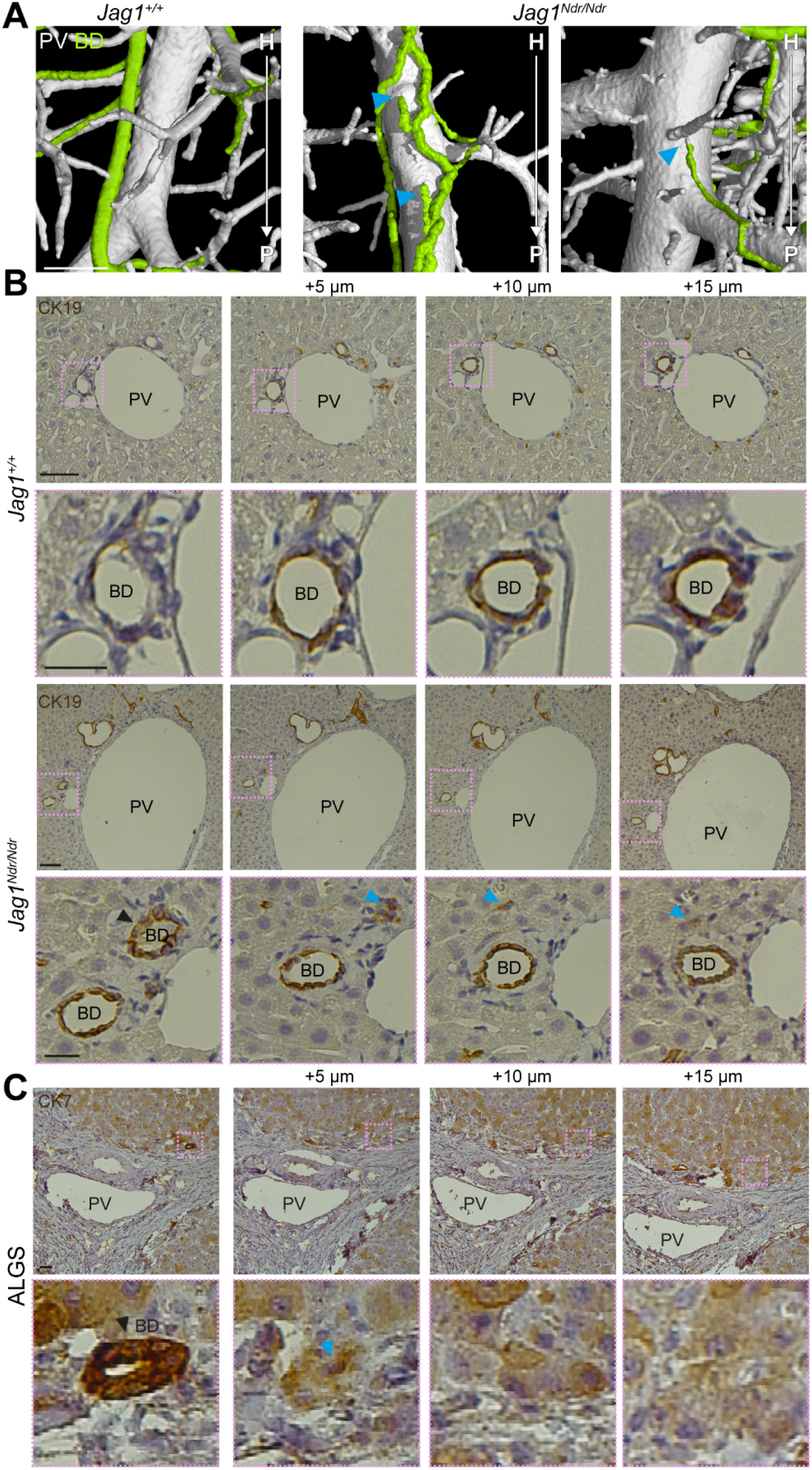
Bile ducts in *Jag1*^*Ndr/Ndr*^ mice and patients with Alagille syndrome terminate abruptly. (A) *Jag1*^*Ndr/Ndr*^ BDs (right side) terminate prematurely (blue arrowheads). (B) 2D histological consecutive liver section confirm blunt end BDs in *Jag1*^*Ndr/Ndr*^ liver. Black arrowhead depicts a lumenized well-formed BD that suddenly terminated in the following sections (blue arrowhead). (C) BDs from patients with ALGS terminate abruptly (blue arrowhead) in consecutive liver histological sections. Scale bars 500 μm, (B top row, C) 50 μm, (B bottom row) 20 μm.

### Wild type bile ducts, but not regenerated bile ducts, are positioned superior to portal vein branches

The biliary tree develops via differentiation of many cholangiocytes around the PV that eventually connect to form a hierarchical network. Which cholangiocytes and ductules will mature or disappear and how this process is regulated is largely unknown. DUCT provides a 3D perspective with full rotation capacity of two biological systems in tandem. This unique feature revealed that *Jag1*^+/+^ BDs run parallel to the PV and are positioned superior to PV as shown in 3D reconstructions (Fig 7A, Supp Fig 10A, C left panel, blue and magenta arrowheads) or in 2D scans (Fig 7B, Supp Fig 10B, D left panel, blue and magenta arrowheads). The typical BD position towards the PV in *Jag1*^+/+^ liver is schematically depicted in Fig 7C. The uniform spatial positioning of the BDs was disrupted in *Jag1*^*Ndr/Ndr*^ livers, and ducts randomly occupied multiple positions along the PV (3D reconstruction Fig 7A, Supp Fig 10 A, C right panel and 2D scan Fig 7B, Supp Fig 10B, D right panel, blue and magenta arrowheads). We performed tile scans from the hilar region of transversally sectioned and stained liver to validate the DUCT finding. In *Jag1*^+/+^ liver, BDs were typically positioned superior to the PV (Fig 7D, image #1, 2, 3, 4, left panel). The *Jag1*^*Ndr/Ndr*^ liver tile scans instead revealed BDs randomly distributed around the PV, regardless of the liver region (Fig 7D, right panel), and the majority of PVs had an inferior BD. Altogether, our data suggest that *Jag1*^*Ndr/Ndr*^ biliary tree regenerates postnatally forming a network of tortuous and distant BDs that branch, end and position stochastically in relation to PV.

**Fig 7.**
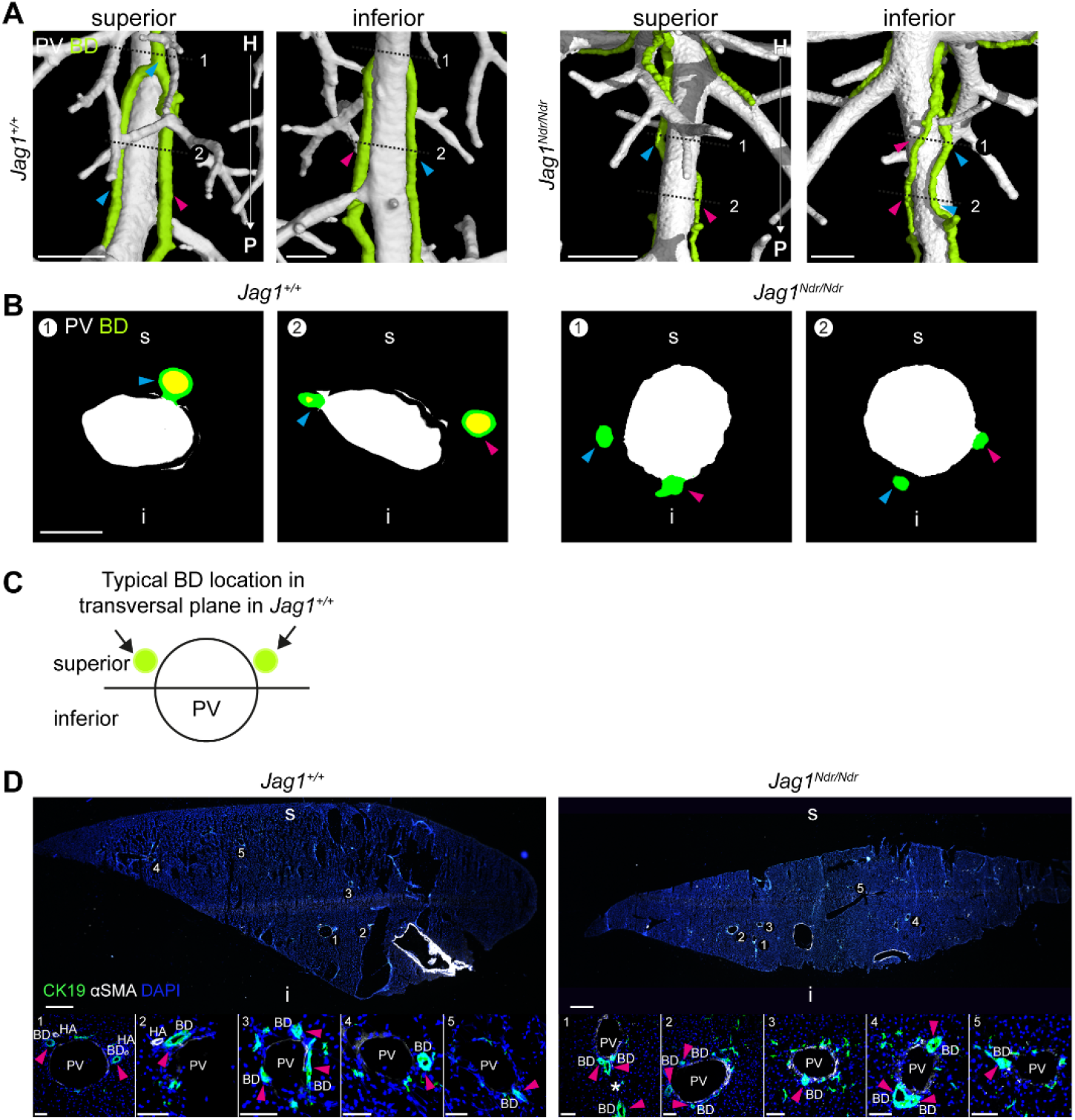
Wild type bile ducts, but not regenerated bile ducts, are positioned superior to portal vein branches. (A) 3D reconstruction of the BD and PV revealed conserved superior positioning of the BD relative to the PV. Scale bar 500 μm. (B) 2D sections of DUCT resin cast delineated in panel A, scale bar 150 μm. (C) Scheme illustrating stereotype BD position in *Jag1*^+/+^ liver. (D) Transverse liver sections confirm the conserved BD position. Scale bars (top panel) 500 μm, (bottom panel) 50 μm.

## DISCUSSION

Precisely defining the three-dimensional architecture of the healthy and diseased liver is a fundamental aspect of liver biology, and improved imaging methods would allow stricter characterization of animal models for human diseases. Until now, 3D analysis has been hampered by a lack of adequate tools and high auto-fluorescence of the tissue. Here, we further improved resin casting to simultaneously visualize, digitalize and quantify the portal venous and biliary system in mouse, and developed a pipeline, using MATLAB and ImageJ, to quantify liver architecture. Using DUCT, we identified several novel phenotypes in the *Jag1*^*Ndr/Ndr*^ mouse model, which we confirmed in liver samples from patients with ALGS (summarized in the Graphical abstract).

Previous resin casting of the biliary system in Notch-deficient liver, lacking hepatic *RBPJК* or *Notch1* and *Notch2* ^29,31^ revealed reduced biliary branching at P120, while constitutive expression of Notch1 intracellular domain (NICD) induced hyper-branching. However at P60, there were no statistically significant differences in biliary branching in *RBPJκ* knockout or NICD gain of function livers ^32^. Similarly, we did not detect significant differences in biliary bifurcations or trifurcations in *Jag1*^*Ndr/Ndr*^ adult mice (4.5 – 6.5 months old) (Supp Fig 9 H – M). However, because DUCT allows analysis of the PV and BD simultaneously, other branching defects were apparent in *Jag1*^*Ndr/Ndr*^ mice, including BD branching in the absence of PV branching, and BD branching occurring distant from PV branch sites (Fig 4). These data suggest that biliary regeneration in the Notch-deficient *Jag1*^*Ndr/Ndr*^ mouse leads to stochastic branching, ignoring portal vein branching, which was also present in patients with ALGS. Interestingly, although there was no difference in the number of branching points per se, the fact that hilar biliary branches were shorter in *Jag1*^*Ndr/Ndr*^ mice (Fig 4) suggests primarily ectopic regenerative branching in the hilar region, rather than in peripheral regions. This would agree generally with the hepatic defects in patients with ALGS, in which bile duct paucity is most prevalent in peripheral liver. Implementing branching quantification as a factor of distance from the hilum would further improve the usefulness of DUCT.

Wrinkling or tortuosity of the biliary system has previously been described for bile-duct ligated rats ^33^ or mice ^34^, and in dogs with chronic liver disease ^35^. In humans, tortuous bile ducts are present in several chronic liver diseases, including hepatitis and cirrhosis ^36,37^. In ALGS, tortuous bile ducts have not yet been reported, but biliary architecture may be an important factor determining liver health. A key study aiming to identify predictors of liver disease outcome ^38^ showed that neither bile duct paucity, nor bile duct proliferation was predictive of outcome. Instead, fibrosis and cholestasis were correlated with outcome, suggesting the functional capacity of the biliary system as a whole, rather than local architecture, dictates outcome. In line with this, a stochastically branching and tortuous biliary system may rescue function, while not leading to a normal density of bile ducts in the periphery.

Previously reported quantification of liver architecture in Notch-compromised mice ^29^ used commercially available software (Scanco) developed for trabecular bone analysis, to analyze biliary volume, branch thickness and number of branch points. DUCT instead allows integrated analysis of the PV and BD system. While DUCT enables 3D analysis of multiple systems simultaneously, the method is not trivial. The resin injections are surgically challenging. Injecting the portal venous system and the biliary system themselves, separately, is more feasible. After one system is injected with resin, the second system is more challenging perhaps due to increased resistance by the presence of resin in the first system. It is also crucial that the resins are fresh, as older resins separate and create artefacts upon microCT scanning. Initially, we aimed to use or mix resins of sufficiently distinct radiopacity to allow automated segmentation and separation of the biliary and vascular systems. However, the green and yellow Microfil resins were not sufficiently distinct at any concentration tested (data not shown) to allow automated separation. Further efforts to test or develop novel resins for automated separation would accelerate the pipeline, which currently relies on manual separation of the portal venous and biliary systems. Finally, the current MATLAB pipeline requires a working memory (RAM) of minimum 32 GB, depending on input data size, and therefore dedicated computers are needed for this analysis.

Identifying and quantifying architectural defects such as branch length, stochastic branching, tortuosity, portal-biliary distance, blunt end bile ducts, or relative orientations of portal venous and biliary system, is very challenging in sections (compare histology and 3D imaging in Fig 3 – 7). We have demonstrated that DUCT is a powerful method for visualization and semi-automated quantitative analysis of two lumenized biological systems in vivo. DUCT has multiple advantages over ink injections and iDISCO, using 3D imaging with microCT, avoiding the drawbacks of tissue autofluorescence or poor antibody penetration. By injecting two resins into a single animal, it is possible to study the relation between the biological systems, which has not been previously reported. While experts in the field are careful to discriminate hilar and peripheral regions of the liver, carefully tracing bile ducts for hundreds of micrometers in sections is not standard practice and is a demanding endeavor. By developing DUCT, it is now possible to map and quantify biliary and vascular architecture in mouse models, setting the stage for an in depth understanding of how these systems interact in health, disease, and regenerative processes.

## Acknowledgements

We thank Kari Huppert and Stacey Huppert for their expertise and help regarding bile duct ligation and their laboratory hospitality. We also thank Nadja Schultz and Charlotte L. Mattsson for their help with common bile duct ligation. We thank Karolinska Biomedicum Imaging Core, especially Shigeaki Kanatani for his help with image analysis. We thank Jan Masek for his scientific input in manuscript writing. The TROMA-III antibody developed by Rolf Kemler was obtained from the Developmental Studies Hybridoma (DSHB) Bank developed under the auspices of NICHD and maintained by The University of Iowa, Department of Biological Sciences, Iowa City, IA52242. We thank Goncalo Brito for illustrating the cal abstract.

