## Supplementary Materials and Methods for "DoUble resin casting micro computed tomography (DUCT) reveals biliary and vascular pathology in a mouse model of Alagille syndrome"

#### MICROFIL® injections

MICROFIL® (Flow Tech Inc.) was prepared as follows. Yellow MICROFIL® (Y) cat. #MV-122 was diluted with clear MICROFIL® (C) cat. # MV-Diluent in 1:3 (Y:C). Blue MICROFIL® (B) cat. #MV-120 was diluted 1:1 (B:C) with clear MICROFIL®. Diluted yellow MICROFIL® was mixed with diluted blue MICROFIL® 1:1 creating a green MICROFIL®. Yellow MICROFIL® was injected into common bile duct (CBD). Green MICROFIL® was injected into portal vein (PV). 1 ml of diluted MICROFIL® is mixed with 50 µl of hardener (supplied by Flow Tech Inc.) prior injection.

Mice were sacrificed by $CO_2$ inhalation and perfused through the heart with Hanks' Balanced Salt solution (HBSS) (Life Technologies cat. # 14025092) for 3 min (perfusion rate 5 ml / 1 min).

Injection into CBD. A small transversal incision was made in inferior vena cava with spring scissor to release the liver vascular pressure. CBD was exposed by moving aside the liver and intestine and cleaned from surrounding tissue in area about 5 mm long. Silk threat (Agnthos AB cat. #14757) was loosely wrapped around the cleaned CBD. A longitudinal CBD incision was made at the spot where CBD enters the pancreas next to sphincter of Oddi by spring scissor. Tubing (PE10, BD Biosciences cat. # 427401) ~ 15 cm long connected to (27G) needle was inserted into the CBD incision. The tubing was prepared by stretching one side of the tube until the diameter becomes thin enough to fit into CBD. Diagonal cut is made at the tip of the tubing while the other side contains needle connected to the syringe filled with MICROFIL®. Yellow MICROFIL® was injected into the CBD until resistance was met or MICROFIL® spots were visible on the liver surface. Massaging the liver with cotton swab while injecting helped to disperse the MICROFIL®. The tubing was removed and silk threat was tightened around the CBD to prevent leakage.

Injection into PV. PV was cleaned from surrounded tissue. A small incision was made in PV using spring scissor. Silk threat was loosely wrapped around the cleaned PV above the incision. Tubing (PE10) ~ 15 cm long connected to (27G – 30G) needle was inserted into the PV incision. Green MICROFIL® was injected into the PV until blood vessels on the surface were filled or resistance was met. Massage the liver with cotton swab while injecting to help disperse the MICROFIL®. The tubing was removed and silk threat was tightened around the PV to prevent leakage.

Liver was dissected out and placed at 4°C overnight (ON) for MICROFIL® to solidify. The next day the liver was fixed with 3.7% formaldehyde solution (FA) (Sigma-Aldrich cat. #F1635) diluted in Dulbecco's phosphate-buffered saline (DPBS) (Life Technologies cat. # 14190144). After 24 hours, liver was washed and kept in DPBS. Liver was separated into lobes. The left lateral lobe was placed in 50% methanol (Sigma-Aldrich, cat. # 322415) for 4 hours and into 100% methanol ON. Further, the lobe was placed in benzyl alcohol (Sigma-Aldrich, cat. #402834) and benzyl benzoate (Sigma-Aldrich, cat. #B6630) (BABB 1:2) solution until transparent. The right medial lobe (only FA fixed) was used for µCT scanning.

### Whole mount immunohistochemistry

Mice were anesthetized by isoflurane inhalation (~2%) and transcardially perfused with HBSS for 3 min (perfusion rate 5 ml / 1 min) and 10% neutral buffered formalin (NBF) for 5 min. Liver was dissected out and further immersion fixed with 10% NBF ON at 4°C. The next day liver was washed and kept in DPBS and separated into lobes. Right medial lobe was stained and cleared following the iDISCO+ protocol and imaged by light sheet microscope by Gubra (Denmark).

Fixed and washed samples were dehydrated in methanol/$H_2O$ gradient: 20%, 40%, 60%, 80% and 2 x 100% methanol, each step 1 hour at room temperature (RT). The samples were bleached in cooled fresh 5% $H_2O_2$ in methanol ON at 4°C. The samples were subsequently rehydrated in methanol/PBS series: 80%, 60%, 40%, 20%, with 0.2% Triton X-100, 1 hour each at RT. They were washed in PBS with 0.2% Triton X-100 (PTx.2) for 2×1 hour at RT.

### Whole organ immunolabelling (iDISCO+)

Samples were incubated in permeabilization solution at 37°C for 3 days. Blocking is carried out in blocking solution at 37°C for 2 days. The samples were incubated with primary antibody in PTwH/5%DMSO/3% donkey serum at 37°C for 7 days. They were washed in PTwH for 1×10 minutes, 1×20 minutes, 1×30 minutes, 1×1 hour, 1× 2 hours and 1× 2 days. Samples were incubated with secondary antibody in PTwH/3% donkey serum at 37°C for 7 days, followed by washes in PTwH: 1×10 minutes, 1×20 minutes, 1×30 minutes, 1×1 hour, 1× 2 hours and 1× 3 days. All steps were performed in tightly closed tubes to minimize evaporation and oxidation.

*Solutions for iDISCO*

PTx.2 (1L): 100ml PBS 10x, 2ml TritonX-100

PTwH (1L): 100 PBS 10x, 2ml Tween-20, 1 ml of 10 mg/ml heparin stock solution

Permeabilization solution (500ml): 400ml PTx.2, 11.5 g glycine, 100 ml DMSO

Blocking solution (50 ml): 42 ml PTx.2, 3 ml donkey serum, 5 ml DMSO

Secondary antibody: Alexa Fluor® 488 (dilution 1:1000, Life technologies)

*Tissue clearing*

Tissue was cleared in methanol/$H_2O$ series: 20%, 40%, 60%, 80% and 100% for 1 hour each at RT. Samples were incubated for 3 hours (with shaking) in 66%DCM (Dichloromethane)/33% methanol at RT and in 100% DCM 15 minutes 2x (with shaking) to remove traces of methanol. They were incubated in DiBenzyl Ether (DBE) (without shaking).

*Light sheet microscopy*

Tissue samples were imaged using a light sheet microscope (UltramicroscopeII, Miltenyi). DBE was used as clearing agent during data acquisition. Data was collected at room temperature using Lavision ultramicroscope system and MV PLAPO 2X C/0.5 objective with dry lens, RI correction collar using Andor Zyla 4.2 Plus sCMOS camera. CK7 staining was detected with AF790 (Alexa Fluor, Life Technologies). The acquisition software used was ImSpector (LaVision biotech).

### Ink Injections

Mice were sacrificed by $CO_2$ inhalation and transcardially perfused with HBSS for 3 min (perfusion rate 5 ml / 1 min).

Injection into CBD and PV. CBD and PV were accessed in the same way as described for MICROFIL® injections. When injecting with ink, there is no need to tighten the CBD with silk threat as the ink is not leaking out. Black ink (Higgins cat. #44032) was injected into the CBD until BDs on the surface were filled or resistance was met. White ink (Higgins cat. #44032) was injected using PE50 tubing (BD Biosciences cat. # 427411, needle size 23 – 25G) into the PV until blood vessels on the surface were filled or resistance was met. Liver was dissected out and separated into lobes. All lobes were cleared in BABB as described above. Liver ink images of right medial lobe were taken under stereomicroscope Stemi 305 (Carl Zeiss Microscopy) using PowerShot S3 IS camera (Canon).

**Liver immunohistochemistry**

5µm FFPE-liver (mouse and human) sections were deparafinized and rehydrated through consecutive baths of xylene (#28975.325, VWR) and isopropanol (#K50655934838, Merck). Endogenous peroxidase was blocked by immersion of the slides in methanol (#322415, Sigma-Aldrich) containing 0,3% $H_2O_2$ (#H1009, Sigma-Aldrich) for 15 minutes and rehydration was finalized by rinsing the slides in tap water. Heat-induced epitope retrieval was done using citrate buffer (PH 6.0) for 20 minutes in a pressure cooker. After blocking of the sections with 2% BSA (#A7906, Sigma-Aldrich) for 20 minutes, slides were incubated for 1 hour at 37°C with primary antibody. Anti-mouse (#G21040, dilution: 1/1000, Invitrogen) or anti-goat (Impress, #MP7405, Vector), respectively, HRP-coupled secondary antibody was applied for 30 min at 37°C and revealed with DAB for 30 seconds (#K3468, Dako). After counterstaining with hematoxylin (#HX86014349, diluted 1/5, Merck), the sections were dehydrated in consecutive baths of ethanol (#20821.310, VWR), isopropanol (#K50655934838, Merck) and xylene (#28975.325, VWR) to finally be mounted with hardening medium (Eukitt, #03989, Sigma-Aldrich).

**Liver Immunofluorescence**

Fresh frozen liver was preserved in Optimal cutting temperature (OCT) compound and placed on dry ice. Fourteen µm sections were briefly fixed with 3.7% FA for 5 minutes and washed 2 x with PBS, 5 minutes. Liver sections were further blocked with blocking buffer (3% Donkey serum in PBS containing 0.3% TritonX-1000) for 1 hour at RT. Primary antibodies were diluted in blocking and incubated ON at 4°C. Liver sections were washed with PBS and stained with secondary antibody (Donkey anti-rat 488, dilution 1:500, Alexa Fluor) and DAPI in blocking buffer for 1 hour at RT. Liver sections were again washed with PBS and mounted using Vectashield (cat. # VEH-1000, Vectorlabs).

**Supplementary table 1. Primary antibody used.**

| | | | |
|---|---|---|---|
| Anti-CK7 | 1:1000 (iDisco) | Abcam | ab181598 |
| Anti-alphaSMA Cy3 | 1:2000 (iDisco), 1:500 (IF) | Sigma-Aldrich | C6198 |
| Anti-CK7 | 1:200 (IHC) | Invitrogen | MA5-11986, clone OV-TL 12/30 |
| Anti-CK19 | 1:50 (IHC), 1:500 (IF) | DSHB | TROMA-III |
| Anti-SOX9 | 1:100 (IHC) | R&D systems | AF3075 |
| DAPI | 1:1000 (IF) | Sigma-Aldrich | D9542 |

*Liver section image acquisition*

The fluorescent images were obtained with LSM880 confocal microscope (Carl Zeiss) with spectral PMT detector using Plan-Neofluar 10X/0.3 and Plan-Neofluar 20X/0.50 objectives at room temperature. CK19 was detected with AF488 (Alexa Fluor, Life Technologies). The acquisition software used was Zen Black (Carl Zeiss).

Chromogenic stained images were taken with AxioImager (Carl Zeiss) microscope, Axiocam 503 color camera using Plan-Apochromat 10x/0.45 M27, Plan-Apochromat 20x/0.8 M27 and Plan-Apochromat 40x/1.4 Oil DIC (UV) VIS-IR M27 objectives at room temperature. The acquisition software used was Zen Blue (Carl Zeiss). Human liver sections were imaged with Nikon Eclipse E1000 (Nikon) microscope, Nikon DS-Fi2 camera (Nikon) using Nikon Plan APO 20x/0.75, Nikon Plan APO 10x/0.40 objectives at room temperature. The acquisition software used was NIS Elements F 4.00.00.

### Image processing

Whole mount liver images cleared with iDISCO+ were initially processed in ImageJ for maximum z-projection and segmentation. The images were filtered using the unsharp mask and integral image filter function. Images were next processed in Amira. In Amira images were filtered using the Gaussian filter and background detection correlation. Images were manually segmented. The manual segmentation was further traced using the autoskeleton function. The skeletons were further analyzed in Amira for length, volume and branching.

Images of ink injected liver were proceeds for filament tracing. Bile duct and portal vein filament tracing was performed using Amira. The images were filtered using the unsharp mask and mean filter. The signal was manually segmented to remove artificial signal. The manual segmentation was further traced using the autoskeleton function. The skeletons were further analyzed in Amira for length, volume and branching. For double ink injection (Figure 1A) the background was changed for esthetic purposes using the lasso tool in Adobe Photoshop.

DUCT 2D slices were exported from MyVGL (Volumegraphics) and processed in ImageJ for maximum contrast and brightness.

Liver sections IHC and IF were processed in ImageJ for better contrast, brightness and fluorescent intensity. Whole liver section tiles were stitched in Zen Black (Carl Zeiss).

### Micro CT measurement

The system GE Phoenix v|tome|x L 240 (GE Sensing & Inspection Technologies GmbH, Germany) equipped with nanofocus X-ray tube (180 kV/15 W) was used for the tomographic measurements that were carried out in the air-conditioned cabinet (fixed temperature 21°C). The samples were adapted for this temperature before the measurement to prevent any thermal expansion effect. To prevent any sample motion during the scanning, the samples were placed in 15 ml Falcon tube, filled with 1% agarose gel. The tomographic reconstruction of acquired data was performed using GE phoenix datos|x 2.0 software. The voxel resolution was fixed for all measurements at 12 µm, except one (sample #2401, 8 µm). Detailed overview of used acquisition parameters is stated in Table 1 in the Supplementary Material.

**Supplementary table 2: Settings parameters of the GE Phoenix v|tome|x L 240 system.**

| Sample | Voxel size | Acceleration Voltage | X-ray tube current | Exposition time | Number of projections |
|---|---|---|---|---|---|
| 2401 | 8 µm | 80 kV | 160 µA | 600 ms* | 2500* |
| 2404 | 12 µm | 80 kV | 160 µA | 600 ms* | 2500* |
| 2405 | 12 µm | 80 kV | 160 µA | 600 ms* | 2500* |
| 2431 | 12 µm | 80 kV | 160 µA | 600 ms* | 2500* |
| 2713 | 12 µm | 80 kV | 160 µA | 334 ms** | 1900** |
| 2714 | 12 µm | 80 kV | 160 µA | 334 ms** | 1900** |

* flat panel DXR250 (2048 px × 2048 px, pixel size 200 µm)

** flat panel dynamic 41|100 ( 4048 px × 4048 px, pixel size 100 µm with binning 2)

*Micro CT data segmentation*

The identification and segmentation of bile duct (BD) and portal vein (PV) canals in each CT cross-section is necessary for further analysis of each system. The segmentation is based on differential contrast between the resin and the soft tissue. Two different resins were used (BD yellow MICROFIL®, PV green MICROFIL®), but the absorption properties of these materials were not distinguishable in CT data.

The resin segmentation was performed by global thresholding and manual corrections using VG Studio MAX 3.2 (Volume Graphics GmbH, Germany) software. The threshold value was determined based on the histogram shape and visual evaluation of a selected cross-section (Supp Fig 11 A, B). The resin cast contained artefacts caused by insufficient filling or high injection pressure. These artefacts include air bubbles in resin, non-homogenous resin contrast (caused by mixing of blue and yellow MICROFIL®) and resin leakage due to lumen rupture (probably caused by high pressure during injection), (Supp Fig 1B). Therefore, the thresholding step was supplemented with manual corrections to create smooth, continuous and solid canal masks. Furthermore, the cut-off for the smallest distal canal included in the mask is considered an area of at least 4 voxels.

Next, the BD and PV systems are identified in the segmented resin mask. In most regions, the BDs blend in with PV in one continuous region (Supp Fig 11 B – D). Manual segmentation is therefore necessary to ensure the correct identification of both systems (Supp Fig 11 C, D). This segmentation was performed by outlining the BD regions in every slice of the CT data. VG Studio automatically creates 3D render based on the regions outlined in CT sections. 3D interactive .PDF files are provided in supplementary for the best assessment of canal complexity (Supp Fig 3 – 8).

*Micro CT data analysis*

All data were analyzed using a custom-written algorithm and freely available Matlab® codes (Version R2017a, The MathWorks Inc., Natick, MA). The DUCT algorithm is designed to analyze morphological parameters of the BD and PV systems, and is compatible with the 3D binary masks. Two separate masks of BD and PV system were generated and the analysis was divided in two independent parts. First, the analysis of the entire portal vein and biliary system and second, analysis of the corresponding main branch (= the longest branch) of each system. For detailed analysis and comparison of whole system versus only the main branch, two algorithms, described by the diagrams in Supp Fig 11 E, F were developed. They differ in the input data and the evaluated parameters. For both algorithms, the first step is to create a 3D skeleton of the input binary mask.

*Skeletonization of DUCT binary masks*

The 3D skeleton was derived using the homotopic thinning algorithm described in [26], specifically optimized for Matlab® implementation by Kollmannsberger [27], (online source: https://www.mathworks.com/matlabcentral/fileexchange/43400-skeleton3d). Calculated 3D medial axis skeleton was subsequently converted to a network graph, using algorithm described in [27], (online source: https://www.mathworks.com/matlabcentral/fileexchange/43527-skel2graph-3d). Resulting network graph is formed by nodes and links between them (Supp. Fig 2 D, G).

*Branching analysis*

Branching points analysis was programmed in Matlab® to analyze the distance between BD and PV branching point. This parameter was calculated using 3D Euclidean distances between the BD branching points and the nearest branching point from PV system. The data is represented as cumulative sum of percentage of BD at a given distances between BD and PV branching points (from 0.015 mm to 3 mm).

For branch length analysis the structure of BD and PV trees were first reconstructed in 3D, using the AnalyzeSkeleton toolbox in ImageJ, which provided the three-dimensional coordinates of all branch points for both BD and PV, as well as the connectivity of the graph. Next, we computed for each branch the length along its path to the euclidian distance between its extremities (branch points). To calculate the generation number of branches (both for the PV and BD structures), we manually defined the origin of the ducts as generation 1, and computed generation number as the number of generation branches separating a given branch from the origin. To distinguish side branching events, we calculated the angle between a branch and its "parent" by computing the dot product p of both their unit vectors. A branch with $p>0.95$ with its parent branch was considered to belong to the same generation. We then computed distributions of length for the BD and PV structures as a function of generation number .The data is represented in two ways; the first is the BD or PV mean branch length in each generation (each branching point) (Supp Figure 9L, M). In the second graph (Figure 4D, 4E), the branching generations are divided into three areas: hilum (either generations 1 – 3, 1 – 5, 1 – 6), intermediate region (either generations 4 – 6, 6 – 9, 7 – 12) and the peripheral region (either generations 7 – 8, 10 – 12, 10 – 13, 13 – 17).

The number of bi-furcations (i.e. one input and two outputs), tri-furcations (i.e. one input and three outputs) and quadri- and more-furcations (one input and more than three outputs), were assessed in Matlab® based on binary mask skeleton nodes that were divided into endpoints and branching points. Branching points closer than 0.2 mm (this threshold value was derived based on visual assessment and knowledge of the system) were merged together and further represented by one node.

*Gap analysis between bile duct and portal vein*

To evaluate the gap between BD and PV the surface distances were calculated in Matlab® for each BD skeleton point by detecting the nearest PV skeleton point and connecting the two points with a line and measuring the non-resin area on this line (zero area in the input binary masks). Surface distance was then calculated using 3D Euclidean distance between the detected non-resin voxel coordinates. The data is represented as cumulative sum of percentage of BD at a given distance from PV (from 0.015 mm to 1.5 mm). The maximum distance between BD and PV for each liver sample is depicted in a separate graph.

*Tortuosity measurements*

To quantify length and tortuosity, total (curved) and theoretical (chord) lengths were measured in Matlab® for the whole system length and for the corresponding main branch. The curved length was defined as a cumulative sum of 3D Euclidean distances between neighboring graph points (i.e. links forming points) multiplied by voxel size. The chord length was defined as cumulative sum of 3D Euclidean distances between neighboring nodes multiplied by voxel size. The chord length therefore reflects system length where any nodes are connected by links with the shortest possible length. To analyze the relationship between BD and PV actual length the BD curved length was divided by PV curved length. Tortuosity was calculated as curved length divided by chord length of the same system. Tortuosity was assessed in %, as BD and PV are not straight lines the actual tortuosity measurements were subtracted by 100% (perfectly straight line).

*Volume analysis*

Total system volume was calculated in Matlab® by multiplying a number of voxels representing PV or BD by volume of one voxel. The relationship between BD and PV volumes was addressed by dividing the BD volume by PV volume.

*Diameter measurements*

The main branch diameter was calculated in Matlab® every 1.5 mm along the total length of the branch. The radius was defined as the minimal distance from the skeleton to the segmented area boundary in the input binary mask (i.e. border between background and area of interest). This boundary was calculated using a two-step procedure. In the first step, the input map was eroded using a 3D spherical shaped structural element with 1 pixel radius. Subsequently the eroded area was subtracted from the original binary mask. This resulted in a binary mask representing the boundary between the background and the area of interest. One radius value at a given skeleton point was then expressed as the minimum distance from that point to the mask boundary. This was calculated, using the minimal value search in the intersection of the boundary mask and the distance map from that point. The distance map from a given skeleton point was calculated as 3D Euclidean distance of the spatial coordinates. Subsequently the diameter value was calculated as the minimum distance to the boundary area multiplied by 2. To avoid any misrepresentation, the one final diameter value at a given point (every 1.5 mm of branch length) was calculated as a mean value of a diameter at that point and diameters at four neighboring points (two on each side). PV and BD diameters were divided into 3 areas: hilum, intermediate and periphery. Hilar region represents distance from 0 – 1.5 (sample #2401) or 0 – 3 mm (other samples), Intermediate region is from 3 – 6 mm (sample #2401) or 4.5 mm – 9 mm (other samples), Periphery is from 7.5 mm – 9 mm (sample #2401) or 10.5 – 13.5 mm (other samples). BD to PV diameter ratio was calculated by dividing BD diameter at a given region by PV dimeter of the same region.
