## Supplementary figures and images for "DoUble resin casting micro computed tomography (DUCT) reveals biliary and vascular pathology in a mouse model of Alagille syndrome"

### Supplementary Figure 1

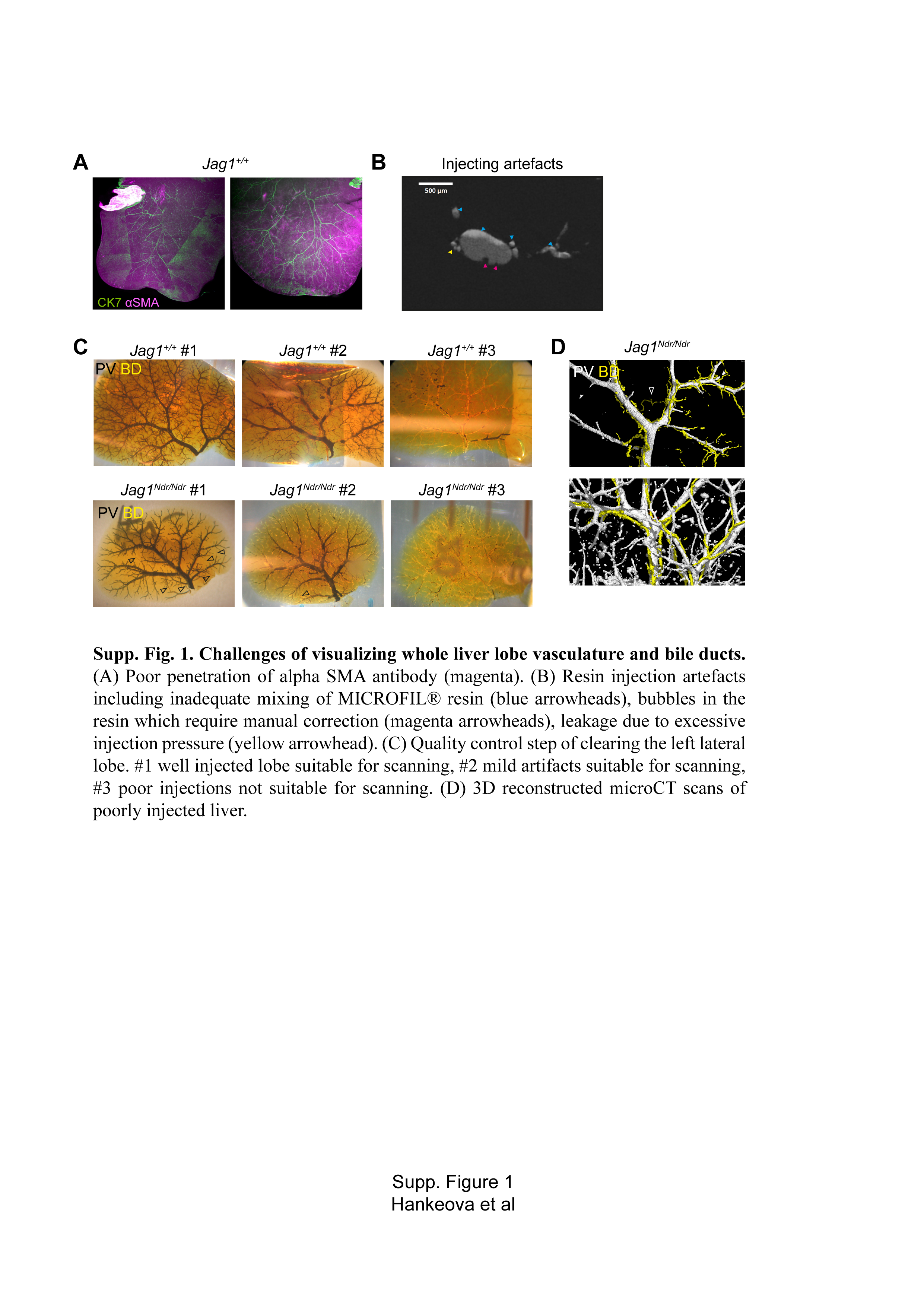

### Supplementary Figure 3

#### Jag1Ndr/Ndr sample 2401

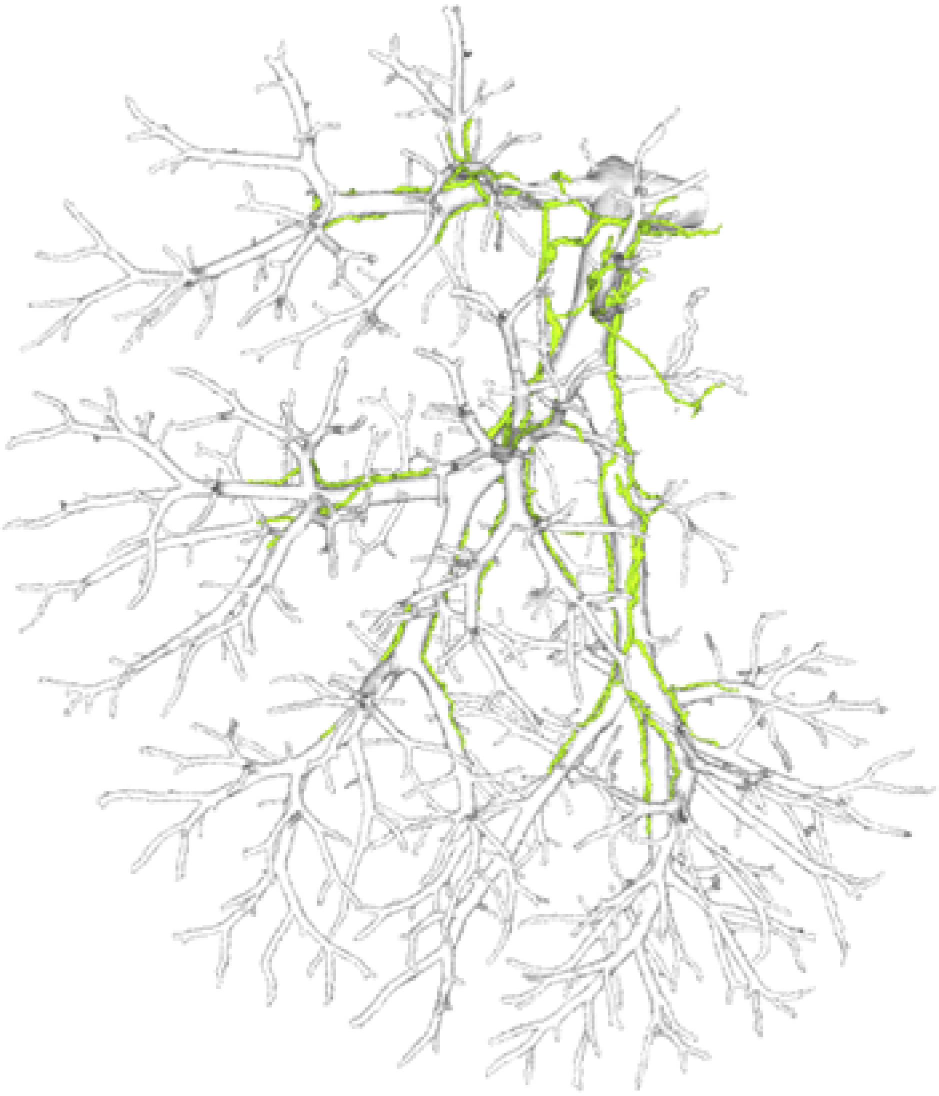

BD+PV

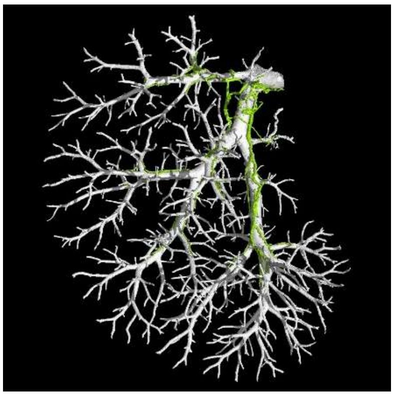

BD

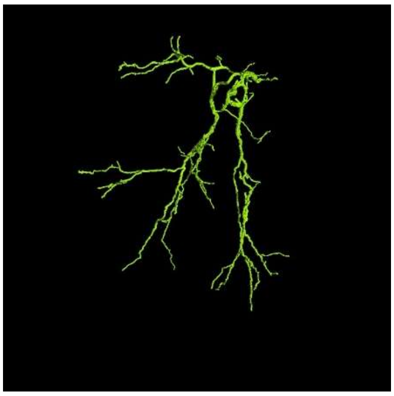

PV

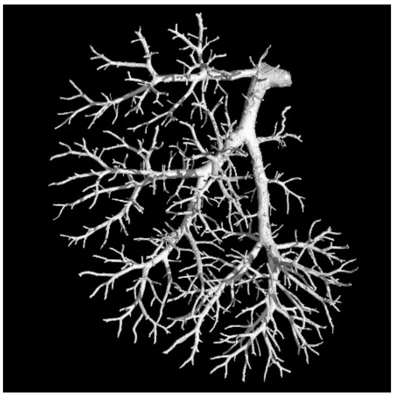

CTLAB
X-ray Computed Tomography

### Supplementary Figure 4

#### Jag1+/+ sample 2404

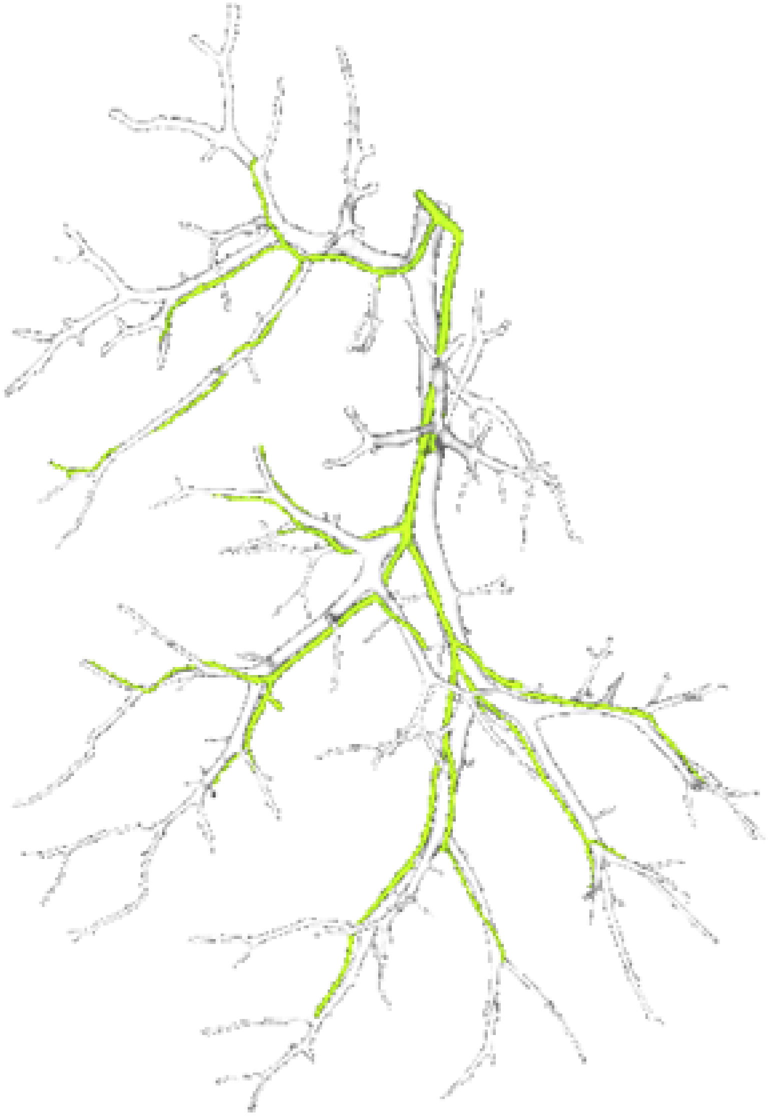

BD+PV

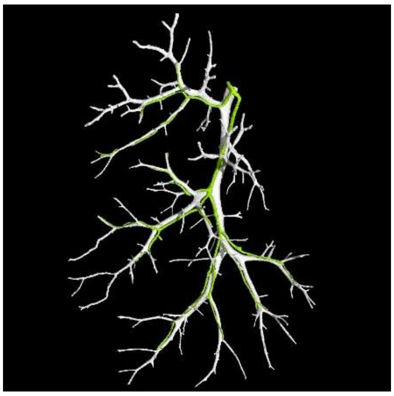

BD

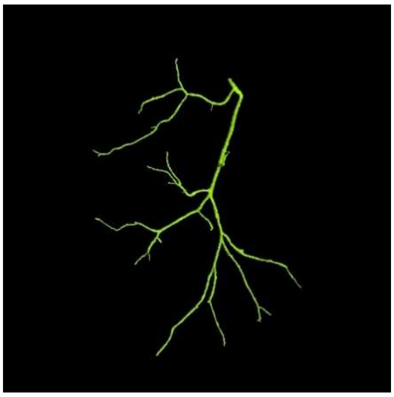

PV

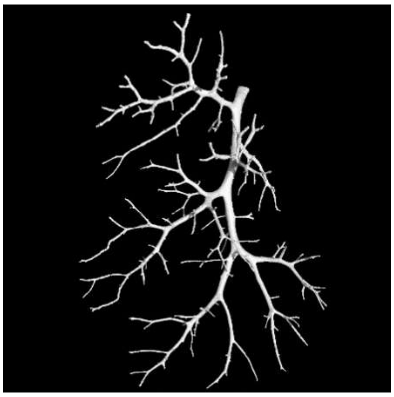

CTLAB
X-ray Computed Tomography

### Supplementary Figure 5

#### Jag1Ndr/Ndr sample 2405

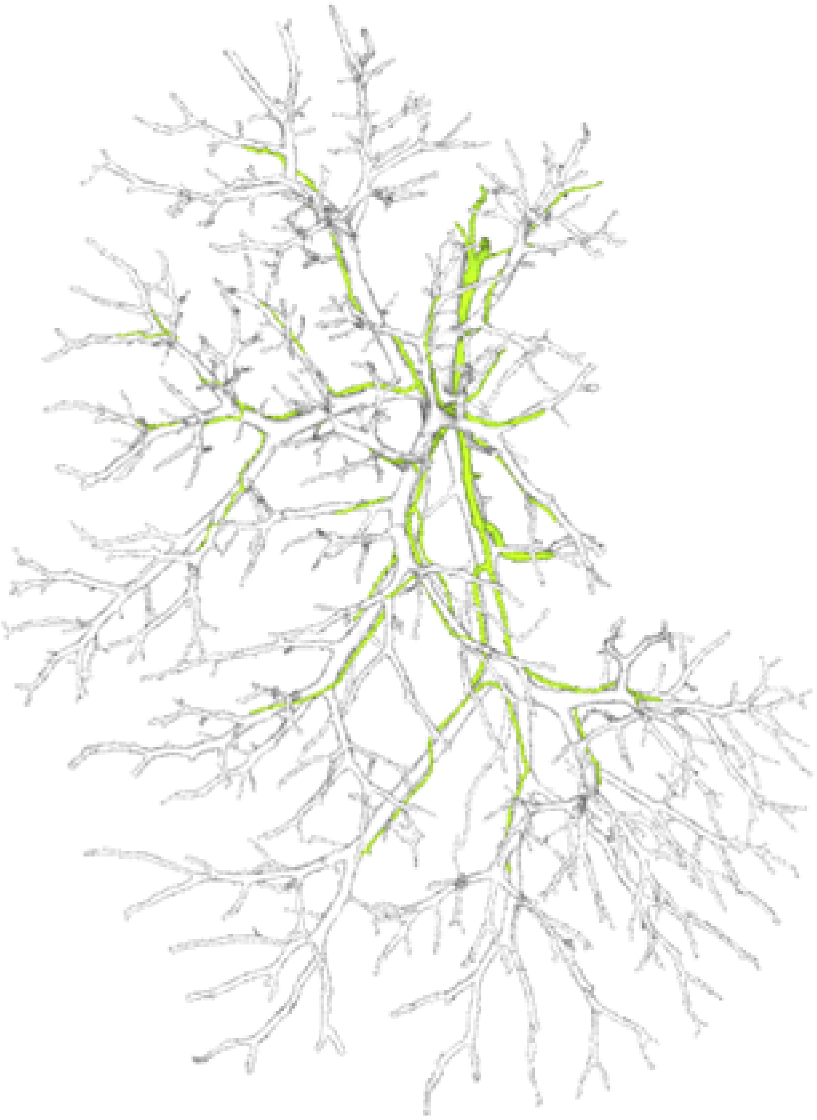

BD+PV

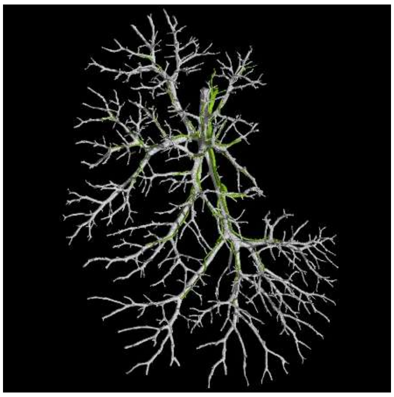

BD

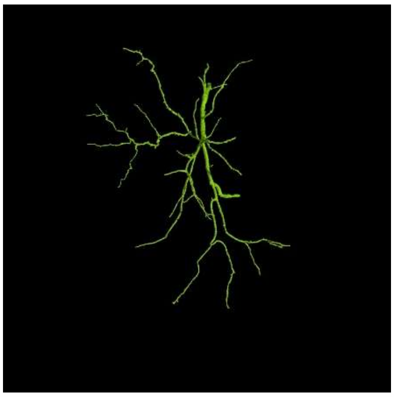

PV

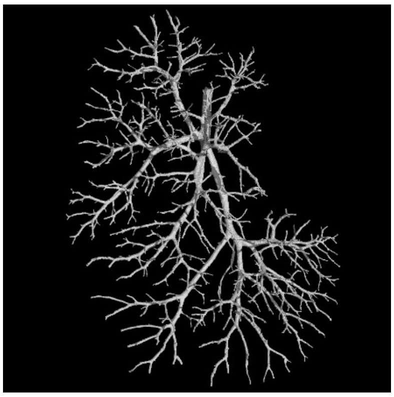

CTLAB
X-ray Computed Tomography

### Supplementary Figure 6

#### Jag1+/+ sample 2431

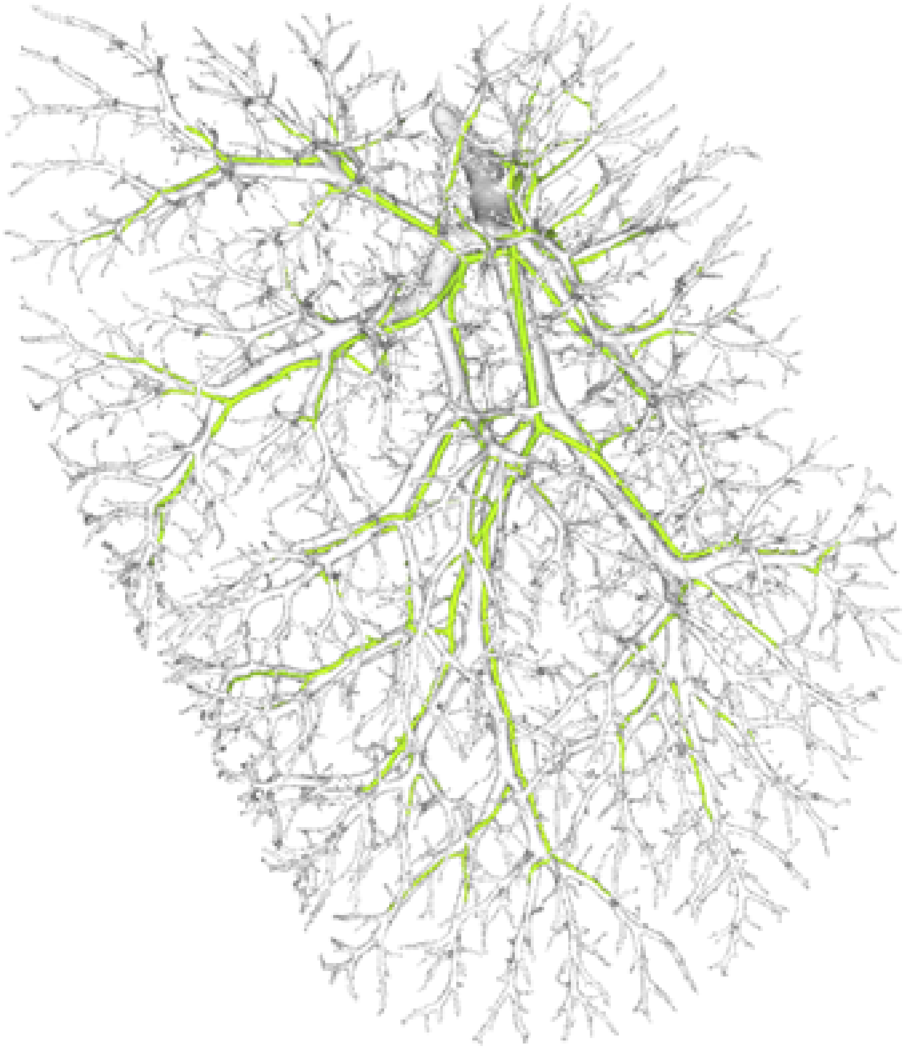

BD+PV

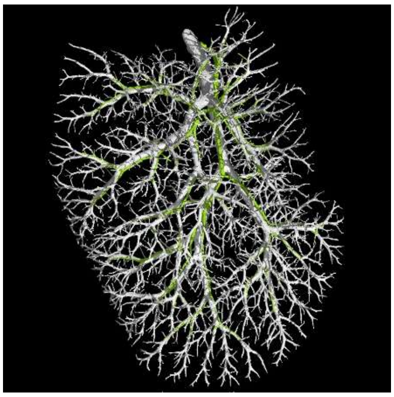

BD

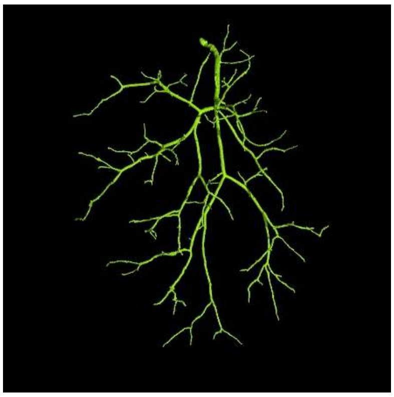

PV

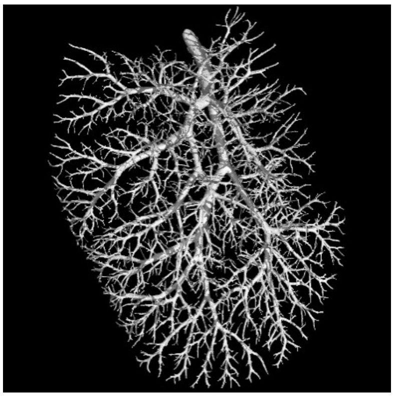

CTLAB
X-ray Computed Tomography

### Supplementary Figure 7

#### Jag1+/+ sample 2713

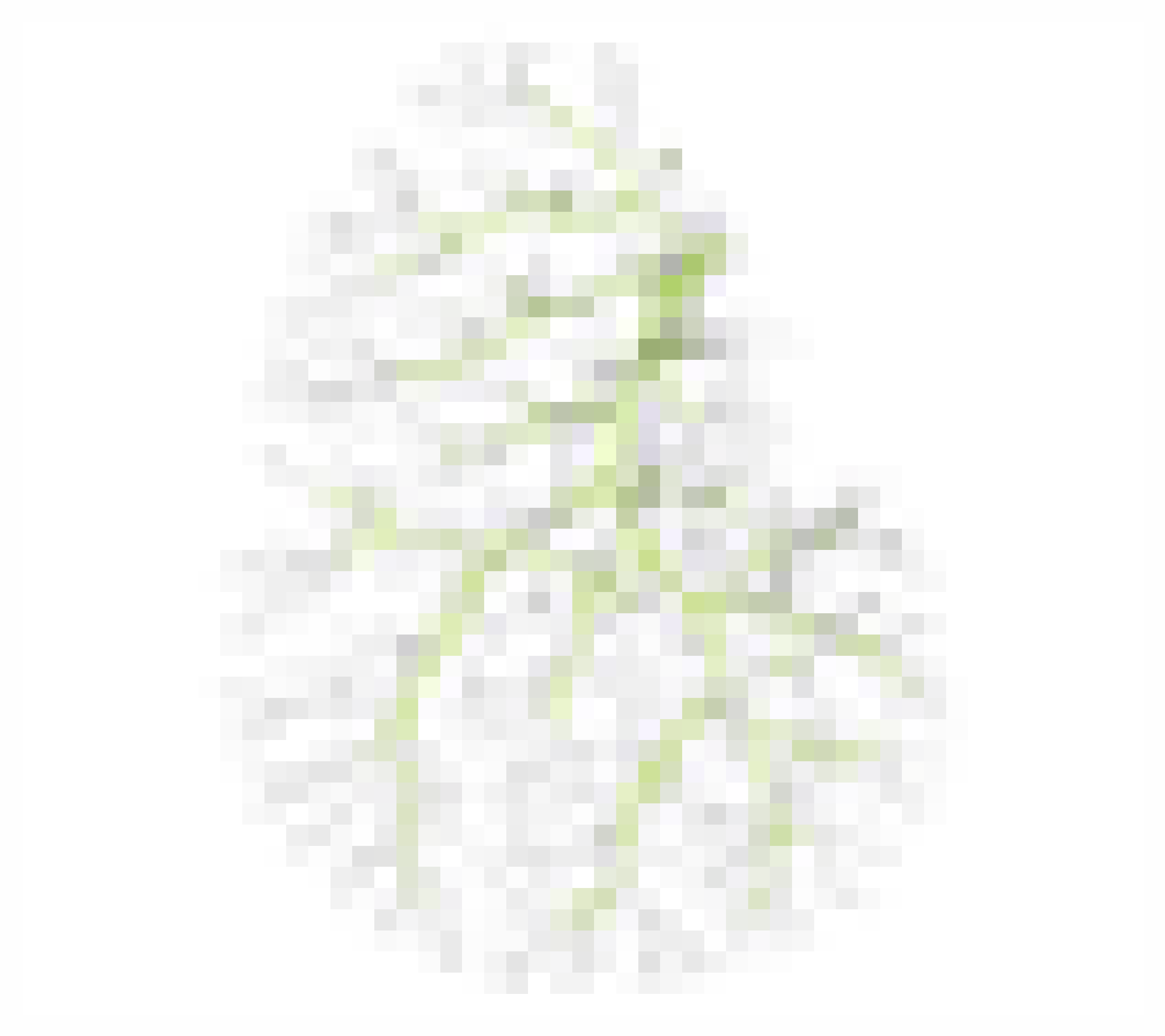

BD+PV

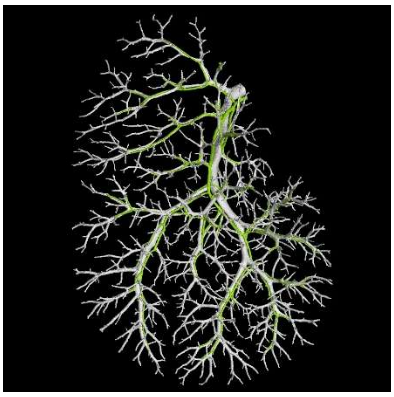

BD

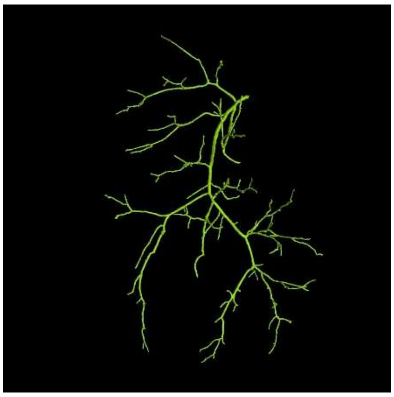

PV

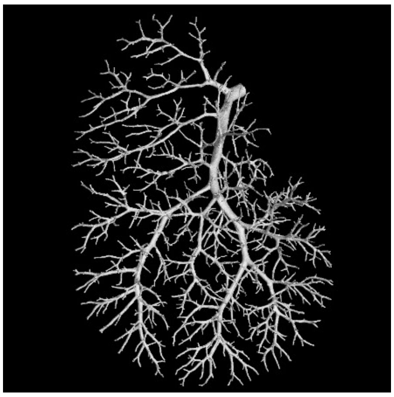

CTLAB
X-ray Computed Tomography

### Supplementary Figure 8

#### Jag1Ndr/Ndr sample 2714

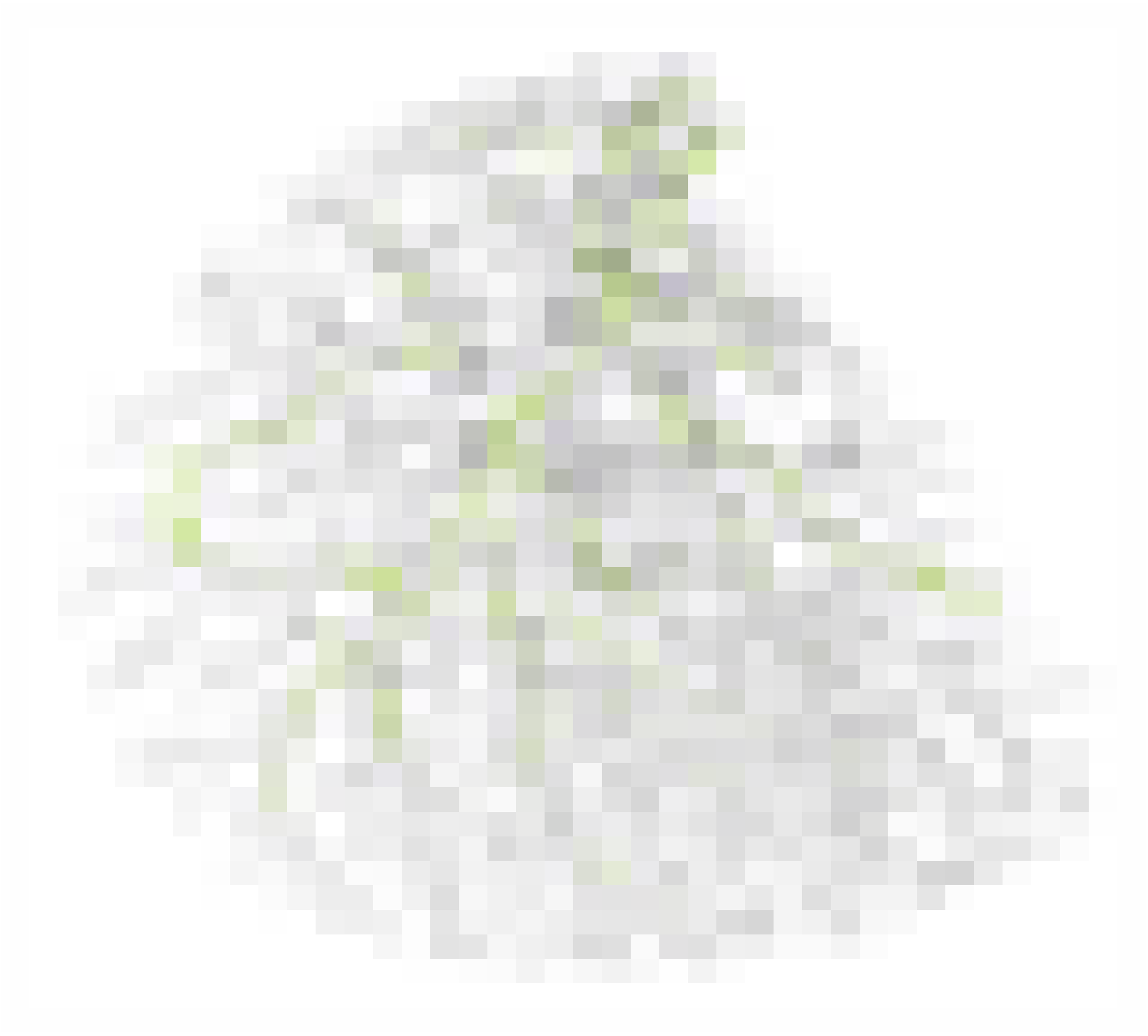

BD+PV

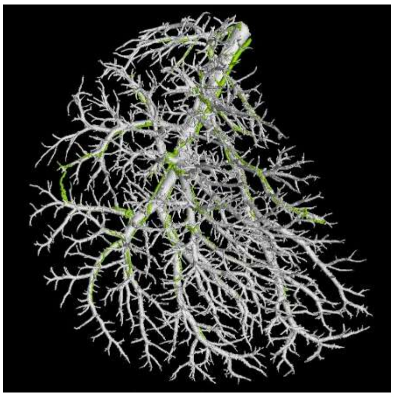

BD

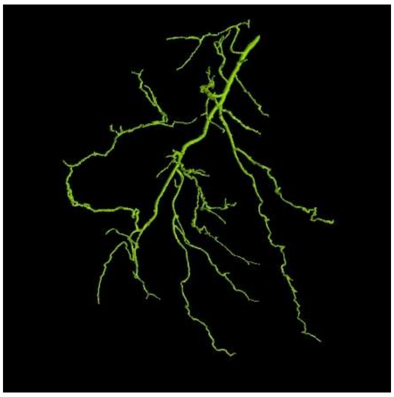

PV

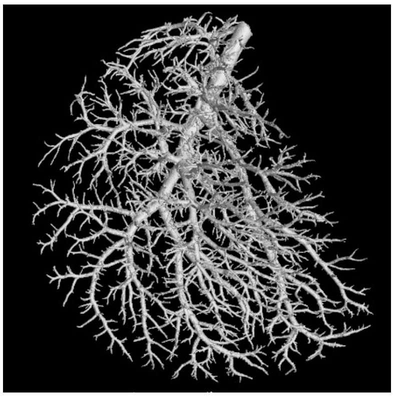

CTLAB
X-ray Computed Tomography

### Supplementary Figure 9

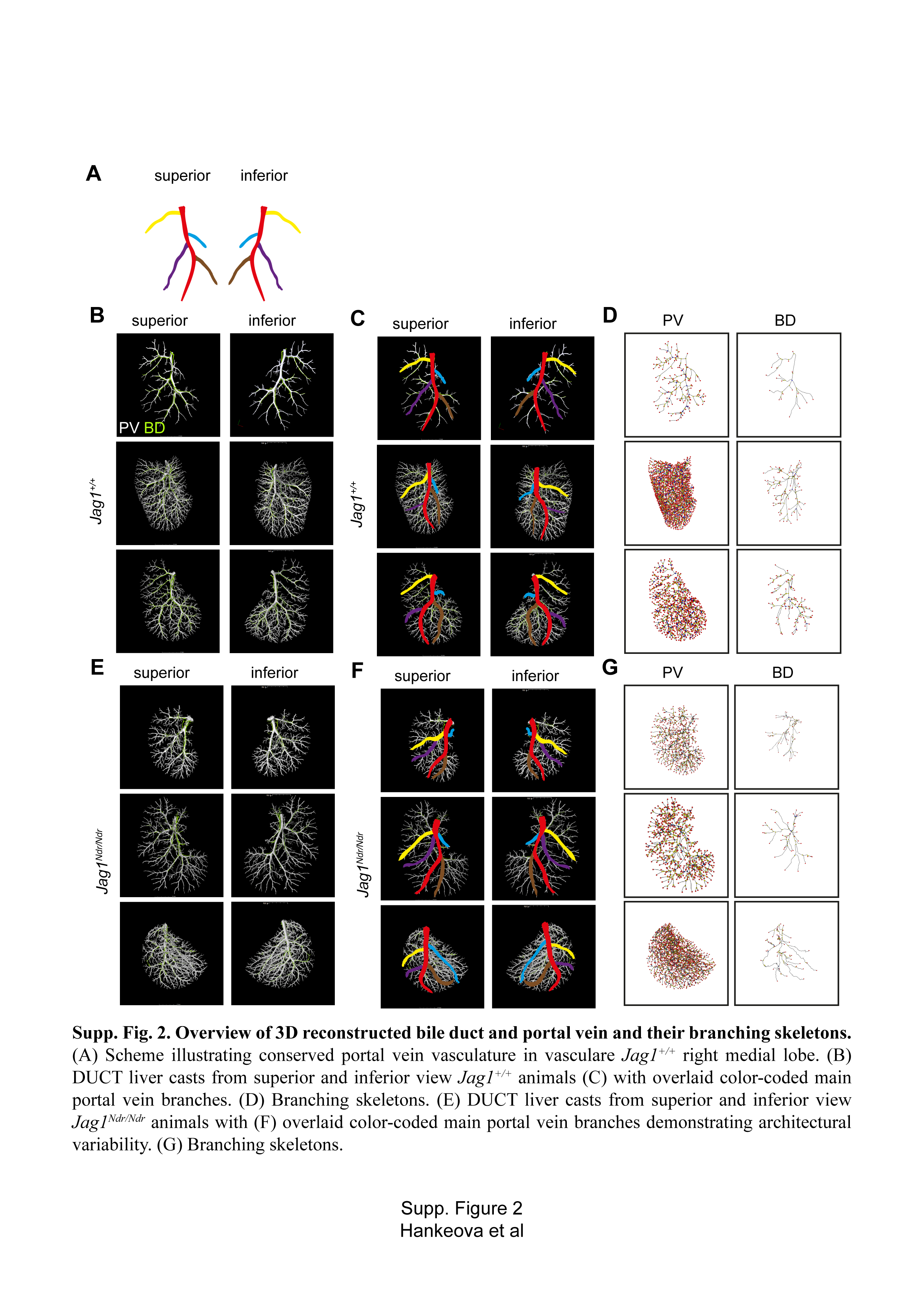

### Supplementary Figure 9

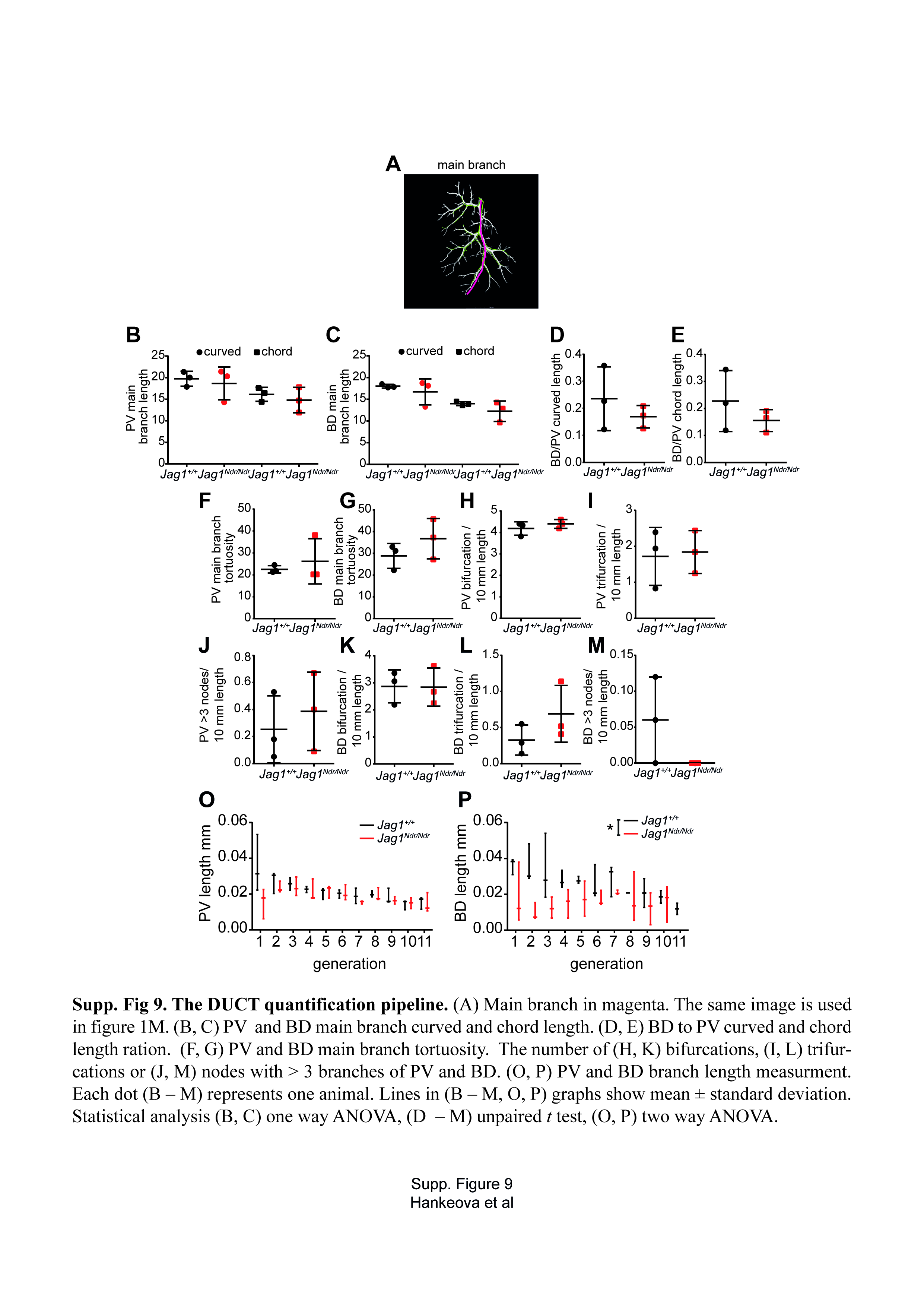

### Supplementary Figure 10

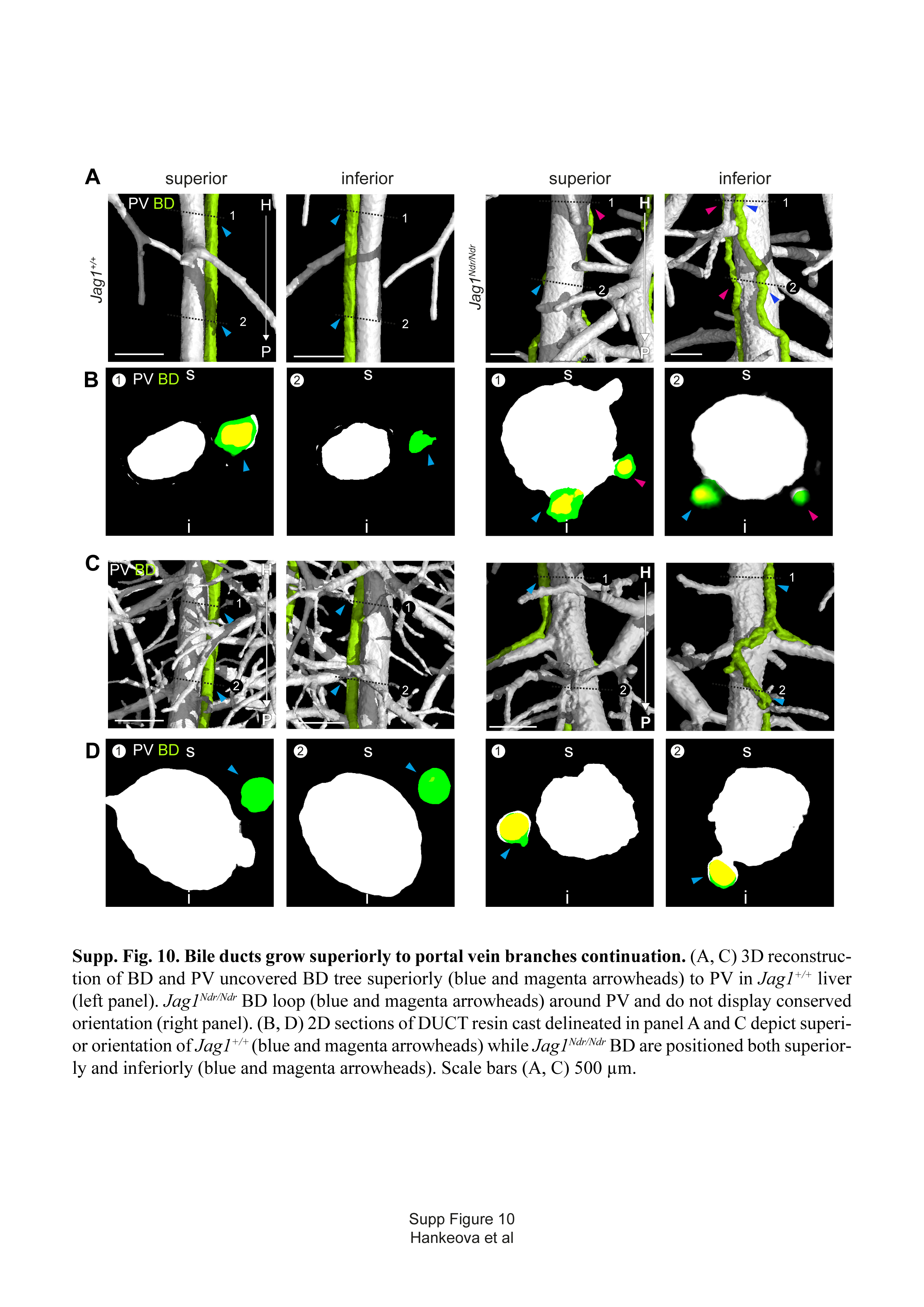

### Supplementary Figure 11

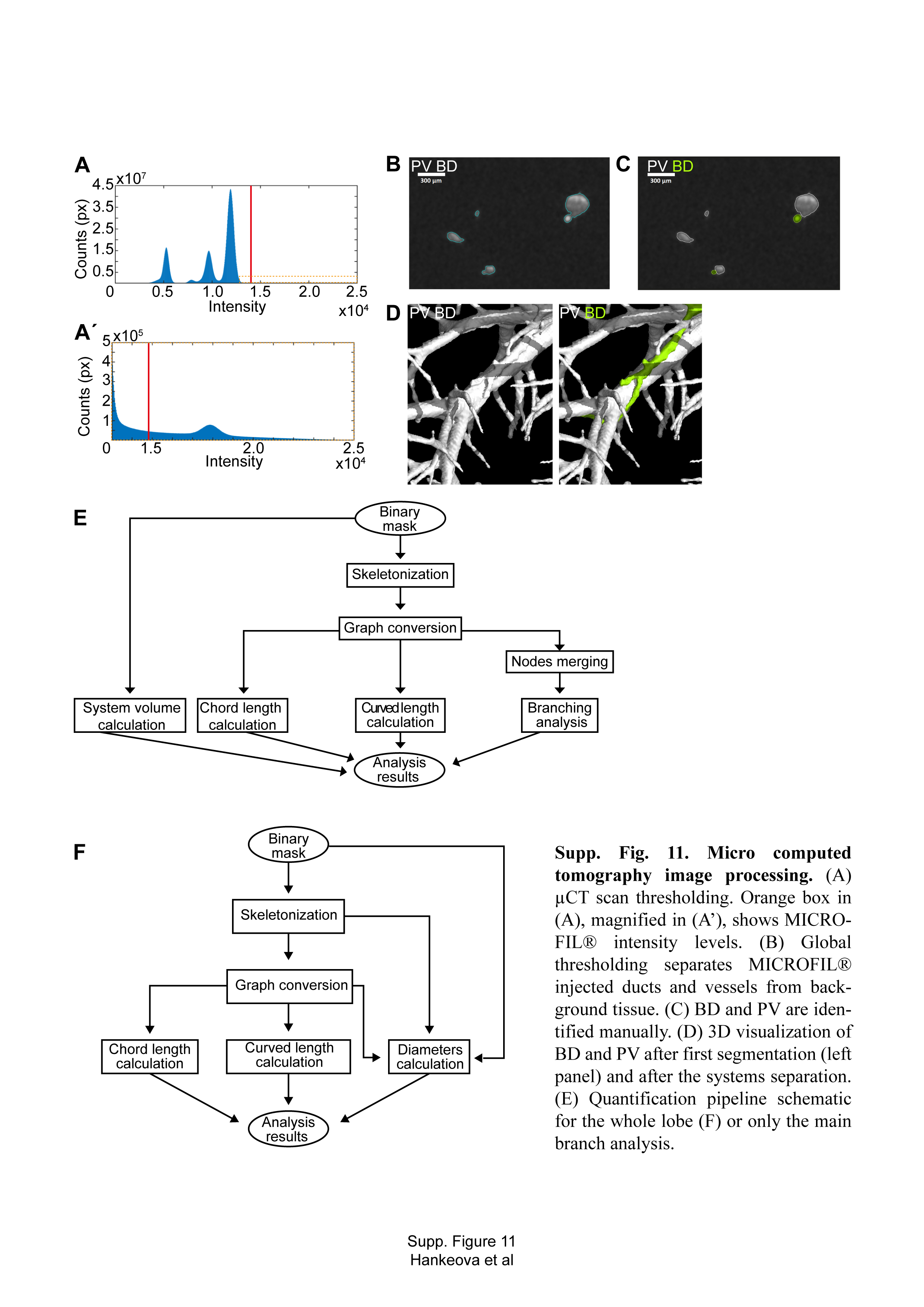
